# CD4^+^ Mucosal-associated Invariant T (MAIT) cells express highly diverse T cell receptors

**DOI:** 10.1101/2025.02.06.636785

**Authors:** Rimanpreet Kaur, Nezar Mehanna, Atul Pradhan, Danielle Xie, Kelin Li, Jeffrey Aubѐ, Barbara Rosati, David Carlson, Charles K. Vorkas

**Affiliations:** Division of Infectious Diseases, Department of Medicine, Renaissance School of Medicine at Stony Brook University, Stony Brook, NY, 11794; Department of Microbiology and Immunology, Stony Brook University, Stony Brook, NY 11794; Center for Infectious Diseases, Stony Brook University, Stony Brook, NY, 11794; Division of Chemical Biology and Medicinal Chemistry, UNC Eshelman School of Pharmacy, University of North Carolina at Chapel Hill, Chapel Hill, NC, 27599; Department of Physiology and Biophysics, Renaissance School of Medicine at Stony Brook University, Stony Brook, NY, 11794; Institute for Advanced Computational Science, Stony Brook University, Stony Brook, NY 11794

**Keywords:** CD4**^+^** MAIT, TCR diversity, CITEseq, CDR3α sequence

## Abstract

Mucosal-associated invariant T cells are highly conserved innate-like T cells in mammals recognized for their high baseline frequency in human blood and cytotoxic effector functions during infectious diseases, autoimmunity, and cancer. While the majority of these cells express a conserved CD8αβ+ TRAV1-2 T cell receptor recognizing microbially-derived Vitamin B2 intermediates presented by the evolutionarily conserved major histocompatibility complex I-related molecule, MR1, there is an emerging appreciation for diverse subsets that may be selected for in humans with distinct functions, including subpopulations that co-express CD4. Prior work has not examined T cell receptor (TCR) heterogeneity in CD4^+^ MAIT cells, largely due to bias of identifying human MAIT cells as CD8^+^ TRAV1-2^+^ cells. In this study, we adopted an unbiased single-cell TCR-sequencing approach of total MR1-5-OP-RU-tetramer-reactive T cells and discovered that CD4^+^ MAIT cells express highly diverse TRAV1-2 negative TCRs. To specifically characterize this TCR repertoire, we analyzed VDJ sequences of single MR1-5-OP-RU tetramer^+^ MAIT cells across two datasets and identified distinct TCR usage among CD4^+^ MAIT cells including TRAV21, TRAV8 (TRAV8-1, TRAV8-2, TRAV8-3), and TRAV12 families (TRAV12-2, TRAV12-3), as well as more variable J chain and CDR3 sequences. Non-TRAV1-2 MAIT cell TCRs were also enriched after in vitro expansion, including with *Mycobacterial tuberculosis*. These results indicate that mature human CD4^+^ MAIT cells adopt distinct TCR usage from the canonical TRAV1-2^+^ CD8^+^ subset and suggest that alternative MR1 ligands in addition to riboflavin intermediates may select them.

## Introduction

Mucosal-associated invariant T (MAIT) cells are innate-like lymphocytes and are among the first responders during infectious or non-infectious inflammatory diseases, including tuberculosis[1–4]. MAIT cells recognize microbial antigens of the Vitamin B2 (riboflavin) pathway bound to the MHC I-related protein 1 (MR1) receptors on antigen-presenting cells (APCs) [5–9]. They are phenotypically and functionally distinct from conventional peptide-specific T lymphocytes, including high surface expression of C-type lectin receptor, CD161, and conserved T cell receptors (TCRs; most are TRAV1-2^+^ in *Homo sapiens*) [10]. The advent of MR1 tetramers[11–15] has enabled the identification of antigen-specific MAIT cells with the most commonly used MR1 tetramers containing the potent activating riboflavin biosynthetic intermediate, 5-(2-oxopro 6-D-ribitylaminouracil (5-OP-RU) (MR1-5OPRU tetramers) and are used to identify MAIT cells by flow cytometry[1, 3, 11, 12, 14, 16]. The biochemical distinction conferring riboflavin intermediate ligand capacity to bind and activate MAIT cells is a ribityl tail which is necessary for MAIT TCR engagement[17] and is highly biased towards canonical MAIT TCR comprised TRAV1-2α chain paired with TRAJ33/12/20 and TRBV6/20 [7, 18]. Variation in TRBV usage has also been reported to confer microbial pathogen specificity[7].

In contrast to mature murine MAIT cells that are largely CD4^-^CD8^-^ (DN) in blood and tissues[2] or non-human primate MAIT cells that are CD8^+^ [19, 20], mature human MAIT cells can co-express CD8 or CD4 molecules like conventional T cells[1, 3]. While CD8 is believed to stabilize MR1 binding to enhance antigen responsiveness analogous to MHC I [21], the role of CD4 co-expression on MAIT cells is less understood and has not been studied in animal models due to their low prevalence. Thus, it is hypothesized that CD4^+^ MAIT cells are selected in human immunity[20]. We and others previously showed that human CD4^+^ MAIT cells adopt distinct phenotypic states from their CD8^+^ counterparts in healthy states including during asymptomatic human *Mycobacterium tuberculosis* exposure and infection[1, 22–24] including differential expression of IL2R, CCR7, TNFRSF4 and TNFRSF13 proteins, indicating selection by alternative co-stimulatory signals. As CD4^+^ MAIT cells are less responsive to in vitro stimulation with canonical riboflavin-derived activating ligands relative to CD8^+^ MAIT cells, it is also hypothesized that they may respond to distinct classes of MR1 ligands from the riboflavin pathway that remain undiscovered[1]. Prior studies of human MAIT cell TCR diversity have focused on CD8 and TRAV1-2 co-expression and elucidation of TRBV gene usage[25, 26]. In this study, we tested the hypothesis that CD4 and CD8 co-expression in MAIT cells correlates with distinct TCR usage by employing unbiased single cell TCR sequencing analysis of unfractionated MR1-5-OP-RU tetramer^+^ MAIT cell subpopulations. We discovered that CD4^+^ MAIT cells express unexpectedly diverse VDJ genes and resembled private TCRs, including significant variation in TCRα, J segment and CDR3α regions compared to CD8^+^ MAIT cells. Unique TRAV1-2^-^ CD4^+^ MAIT cells TCRs were also detected after in vitro incubation with cytokines and *Mycobacterium tuberculosis* lysates. Together, our results suggest that human CD4^+^ MAIT cell TCRs may be selected by alternative MR1 ligands in addition to canonical riboflavin intermediates.

## Methods

### Human subject recruitment and PBMC isolation

Healthy donors were recruited at Stony Brook University (SBU) under IRB2021-00478 or purchased from the New York Blood Center (NYBC) for use in single-cell transcriptomic and flow cytometric experiments. PBMCs were isolated using Ficoll (Fisher Scientific) or Sepmate tubes (Stem Cell Technologies) using the manufacturer protocol. The isolated PBMCs were stored in Bambanker serum-free media (GC-Lymphotec) in liquid nitrogen and thawed as needed for immunology assays.

### Flow cytometric PBMC stimulation assay

Cryopreserved healthy donor PBMCs (n=13) were thawed and 200,000 cells were cultured for 7 days +/- stimulation conditions detailed below using complete media: RPMI 1640 (Quality Biologicals) supplemented with 10% (v/v) heat-inactivated Fetal Bovine Serum (FBS), penicillin/streptomycin (100 U/ml), L-Glutamine (2 mM), sodium pyruvate (1 mM), non-essential amino acids (0.1 mM), HEPES buffer (10 mM) and 2-mercaptoethanol (2-ME) (50 uM) supplemented with IL-2 (50 U/ml; PeproTech) at 37°C, 5% CO2 in U-bottom 96-well plates. Stimulation conditions included *Mycobacterium tuberculosis* (Mtb) Erdman whole cell lysate (1:100), riboflavin biosynthesis intermediate 5-OP-RU (2mM), folate catabolites Ac-6FP (2uM) and 6-FP (2uM), and non-specific T cell stimulation with anti-CD3/CD28 magnetic beads at a 1:100 bead to cell ratio (Dynabeads; Invitrogen). *Mtb* whole cell lysate was prepared from freshly cultured *Mycobacterium tuberculosis* Erdman culture, OD600-0.8 in 7H9 liquid media in 50 mL prior to bead-mediated lysis in a Biosafety level 3 facility at SBU. MAIT cells were identified as Live CD3^+^MR1-5-OP-RU tetramer^+^CD161^++^ MAIT cells (gating strategy **Fig 1A**).

**Figure 1.**
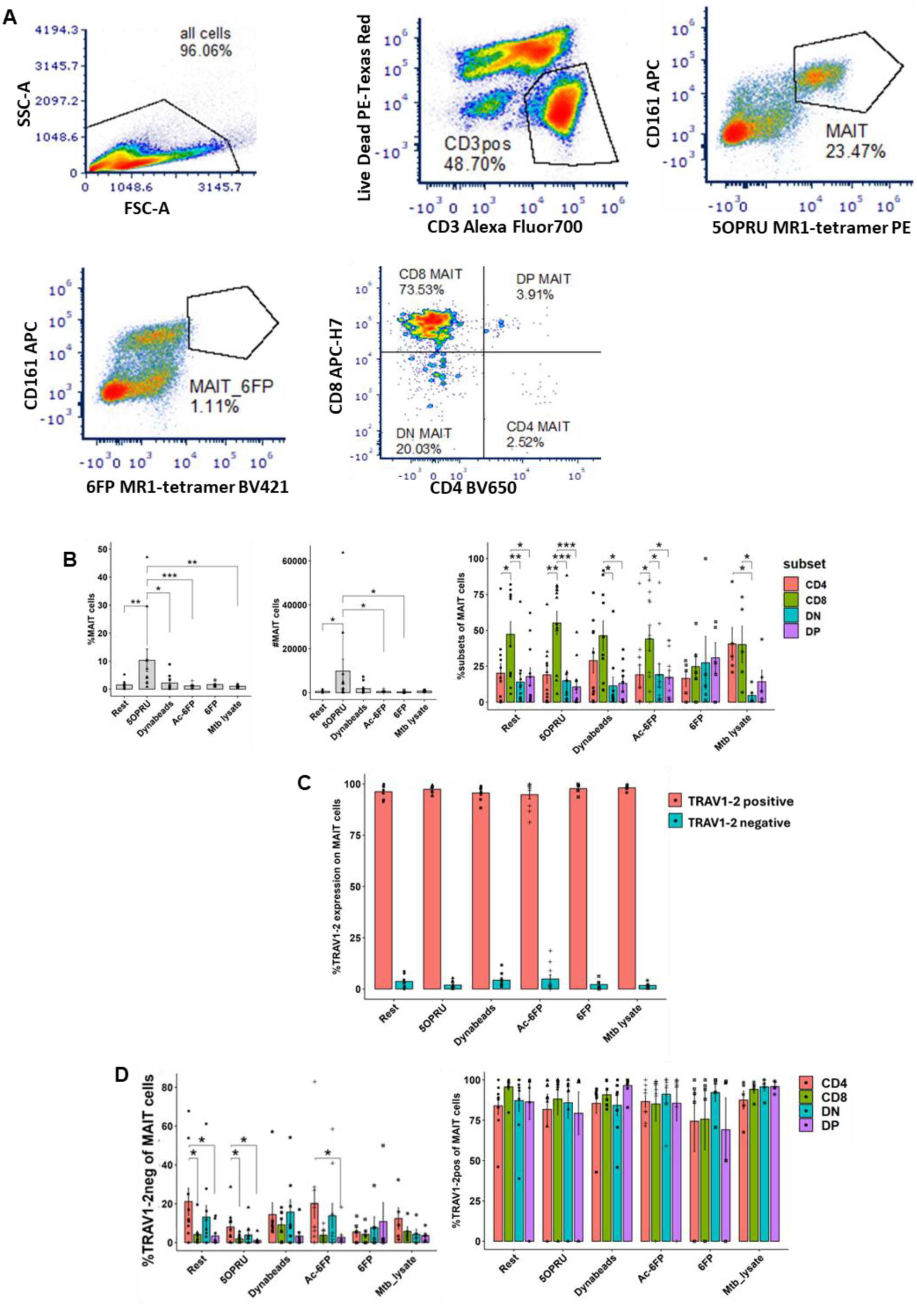
CD4+ MAIT cells are more likely to express TRAV1-2 negative TCRs. **A.** Representative flow cytometric gating strategy of MAIT cells and CD4+/CD8+ subsets after 7 days of in vitro incubation with 5-OP-RU. **B.** Total MAIT cells and subset frequency after 7 days of incubation in vitro with various stimuli. **C.** MAIT cell frequency stratified by TRAV1-2 expression. **D.** Frequency of TRAV1-2 negative (left) and TRAV1-2 positive (right) MAIT cells stratified by subset. Statistical comparisons made by unpaired t-test. *p<0.05

### CITE-seq and VDJ sequencing

Cryopreserved PBMCs from two healthy donors were thawed and sorted directly ex vivo for CITE-seq or cultured in complete RPMI 1640 media in the presence of IL-2 (50U/ml; PeproTech) in a flat non-treated sterile 24-well plate (1M cells/well) with or without *Mtb* whole cell lysate for 7 days at 37°C with 5% CO2 to activate PBMCs.

### Flow cytometry staining and analysis

PBMCs were FcR-blocked (1:20 dilution, BD Pharmingen) for 15 min at room temperature (RT) and stained for cell surface receptors with fluorescent antibody cocktail for 15 mins at RT, followed with IC-fixation (Invitrogen), cells were acquired on Aurora spectral analyzer (Cytek). See Supplemental **Table 1** for complete list of reagents. The flow cytometry analysis was performed on FCS Express 7(De Novo Software), with manual gating, and plotted using ggplot2, rstatix, and dplyr package on Rstudio.

### Single-cell sorting and CITEseq staining

PBMCs were prepared for CITE-seq by FcR-block (ebioscience) for 15min at RT and stained with fluorescent antibody cocktail for cell sorting/enrichment of MAIT cells (CD3^+^, CD161^++^, 5-OP-RU MR1 tetramer^+^), NK cells (CD3^-^, CD14^-^, CD19^-^, CD16^+^CD56^+/-^), TCR γδ cells (CD3^+^, TCR γδ+) and iNKT (CD3^+^, CD161^+^, Vα24Jα18^+^) for 15 mins at RT using Aria FACS in the Stony Brook University (SBU) Flow Cytometry Core Facility. See Supplementary **Table 1** for complete list of reagents. The sorted cells were collected in complete-RPMI media and washed with cold PBS, blocked with Human Truestain FcX (Biolegend) for 10 minutes at 4°C, then stained with hashtag and Total-seqC antibody cocktail for 30min at 4°C (Supplementary Table 1). The cells were pelleted down at 400 rcf for 5 minutes to remove the residual hashtag and Total-seqC antibody cocktail. The cells were resuspended in sterile cold PBS + 0.04% BSA for downstream 5’ single-cell immune profiling assay.

### Single-cell Immune Profiling Assay and Analysis

The cell viability and count of each sample was measured by Countess 3FL with trypan blue staining. Approximately 30,000 cells (5000 cells per sample) were loaded on a 10X Genomics Next GEM Single Cell 5’ Gel Bead Kit in the SBU Single-cell Genomics Core Facility. Gene expression (GEX), Surface Antigen (SA) and VDJ (Tcell-receptor (TCR) libraries were prepared using Chromium Next GEM 5’ Single Cell with Feature Barcode and VDJ assay, according to the manufacturer’s instructions. The libraries were sequenced at a depth of approximately 25,000 (GEX), 10,000 (SA) and 10,000 (VDJ) paired reads per cell on an Illumina NovaSeqX sequencer (Novogene, Inc). The gene expression, surface feature barcode raw FASTQ files were analyzed on Cell Ranger (10X genomics, v.7.0) using multi-pipeline with human reference transcriptome (GRCh38) downloaded from 10x Genomics website and hashtag configuration file, which contained the barcodes related to each sample. The obtained BAM files were converted to FASTQ files and again run on Cell Ranger for SA and VDJ analysis.

The VDJ output was merged with the gene expression matrix using djvdj package on R (v.4.3.2) and then pre-processed using the *Seurat* (v.4.0) package. Quality check was performed where cells with more than 6000 gene counts and more than 5% mitochondrial content were removed. The top 3000 variable features were selected using *FindVariableFunction()* and normalized individual samples before performing integration with the *Harmony* R package. After integrating all samples, principal component analysis was performed and the first 20 principal components were selected for *FindCluster*() with a resolution of 0.8 and visualized with UMAP using default Louvain algorithm. In VDJ data, only cells with paired α and β chain TCRs were selected for further analysis. MAIT cells with one α and one β chain transcript expressed were further stratified by TRAV1-2 positive transcript expression to analyze alternative TCR usage. Single cells with dual TCR α chains were also analyzed.

### Validation of MAIT single-cell transcriptome in a separate data set

To augment the sample size of single MAIT cells and validate our results, we also used a publicly available single-cell transcriptome and VDJ raw dataset generated by CITE-seq of MAIT cells that were similarly defined as MR1-5-OP-RU tetramer^+^ in *Garner et al* annotated experiment #1 [25]. The data were retrieved and analyzed using an earlier version of the pipeline in Rstudio with Seurat package. The cells with more than 8 percent mitochondrial gene content and gene count above 3,000 were removed. Scaling and log normalization were performed to preprocess data and after principal component analysis, the top 20 principal components were chosen for clustering and visualization of cells in UMAP or TSNE. Similarly, VDJ data was combined with single-cell transcriptome using the djvdj package and truly paired TCR with single α and β chains were selected for further analysis.

### GO annotations and TCR functional analysis

The epitope prediction against TRAV1-2^+/-^ MAIT cells was accomplished using McPAS-TCR (Pathology-associated TCR database) (http://friedmanlab.weizmann.ac.il/McPAS-TCR/). This database comprises TCR sequences associated with different pathological conditions and can be queried by TCR composition, CDR3 sequence, epitope, tissue, MHC restriction, and other criteria [27].The CDR3α sequences detected in our study and present in the McPAS-TCR dataset were then used to predict possible epitopes in different pathologies with +/-2 amino acid insertions or deletions. Further, the functional or GO annotations were performed on TRAV1-2 negative and positive MAIT cells to look for their biological functions using KEGG analysis in R and string networking analysis (https://string-db.org/), for which we set a threshold of avg_log2FC above 0.5.

## Results

### CD4^+^ MAIT cells are enriched with TRAV1-2 negative TCRs

To identify the proportion of MAIT cells with non-canonical TCR expression and define their reactivity to MR1 ligands, we first stained healthy donor PBMCs and analyzed MR1-5-OP-RU tetramer^+^CD161^++^ cell frequency in healthy donors (n= 13) using spectral flow cytometry. Analyses were stratified by CD4 or CD8 co-expression resulting in four subsets based on surface expression of co-receptors: CD8^+^, CD4^+^, DN (CD8^-^CD4^-^) and DP (CD8^+^, CD4^+^) (**Fig 1A)**.

We assessed the frequency of each subset at baseline and after in vitro expansion using MR1 ligands or mitogen and observed total MAIT cells expanded with riboflavin derivative, 5-OP-RU, but not with folate derivatives (6FP and Ac-6FP), non-specific T cell stimulation with anti-CD3/CD28 (Dynabeads) or *Mtb* whole cell lysates (**Fig 1B**). We confirmed our previous findings that predominantly CD8^+^ MAIT cells expanded with 5-OP-RU antigen relative to CD4^+^ MAIT cells [1]. We did not observe any significant effect of folate catabolites Ac-6FP or 6-FP on the relative distribution of MAIT cell subsets (**Fig 1B**). Next, we stratified our analyses by TRAV1-2 surface staining and found that most 5-OP-RU-induced MAIT cells were TRAV1-2^+^ (**Fig 1C**) and that the TRAV1-2^-^ MAIT cells were more likely to co-express CD4 than CD8(**Fig 1D, Supplemental Fig 1A-C)**.

### CITEseq defines functionally distinct CD4^+^ and CD8^+^ MAIT cell populations

To define the transcriptional signature and alternative TCR usage of CD4^+^ and CD8^+^ MAIT cell subpopulations, we performed CITE-seq with single-cell VDJ sequencing of sorted MAIT cells (CD3^+^MR1-5-OP-RU tetramer^+^CD161^++^ from two healthy donors directly ex vivo after thaw and after in-vitro incubation for 7 days with IL2 +/- *Mtb* whole cell lysate to select for *Mtb*-reactive MAIT cells. First, we defined the transcriptional and surface barcode antibody profile of all sequenced cells and performed differential gene expression analysis between different cell types (**Fig 2A, B**). The presence of *CD3D*, *CD3G* and *CD3E* gene transcripts, CD3 surface protein expression and differential expression of canonical genes *KLRB1*, *ZBTB16*, *IL18R1*, *CXCR6*, *SLC4A10*, and *TRAV1-2* resulted in four clusters (C1-4) distinguishing MAIT cells from other innate lymphocytes sorted in the same batch (**Fig 2B-D; Supplemental Fig 2**) [28].

**Figure 2:**
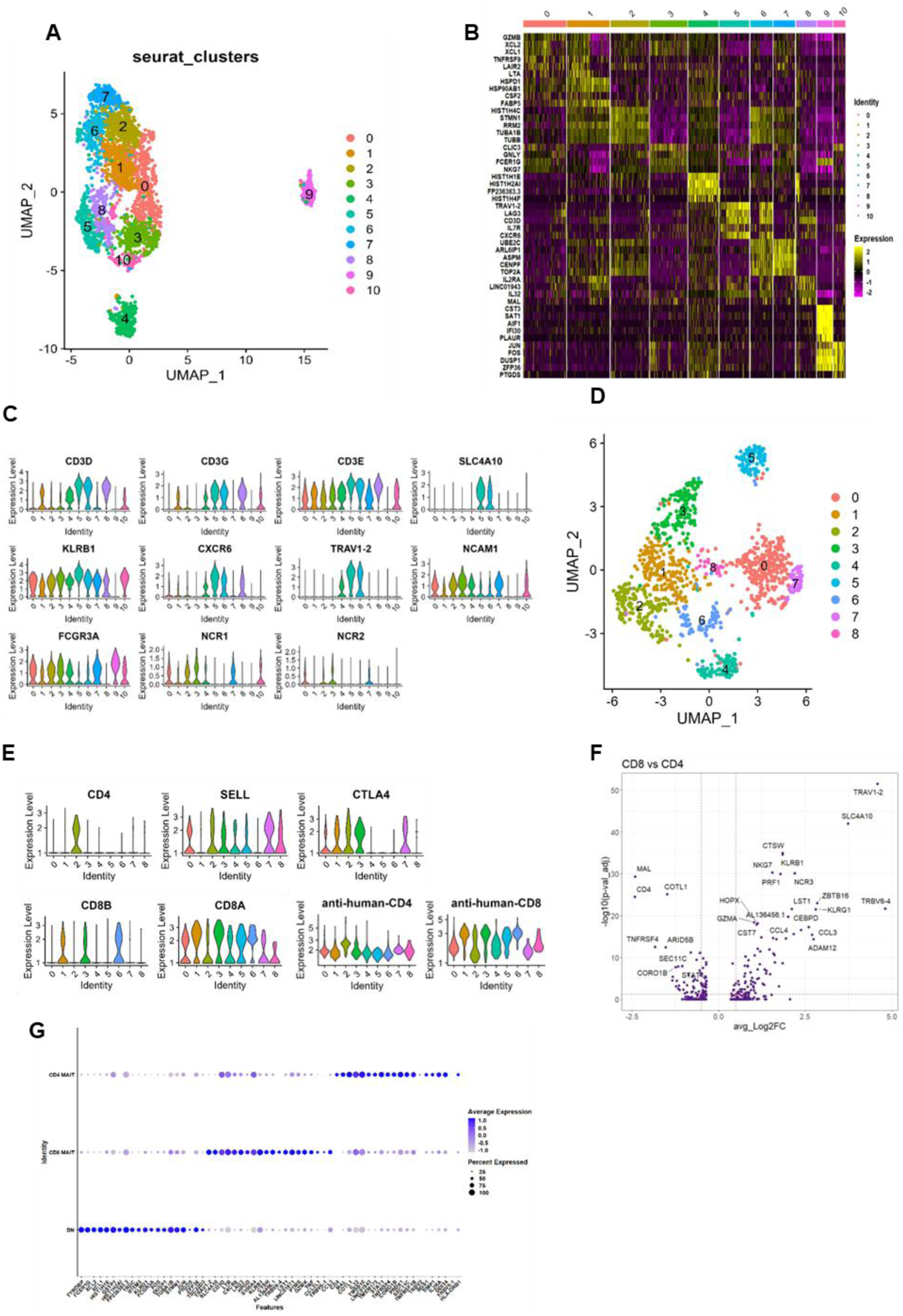
MAIT cell identification and visualization by cellular indexing of transcriptomes and epitopes (CITE-seq) reveals distinct CD4+ and CD8+ subsets. **A.** Uniform Manifold Approximation and Project (UMAP) visualization of all sequenced cells. **B.** Heat map displaying the top differentially expressed genes in Seurat clusters. **C.** Violin plots demonstrating differential gene expression analysis between clusters. **D.** UMAP visualization of MAIT cell subclusters. **E.** Violin plots displaying the expression of CD4 and CD8 genes in MAIT cell clusters. **F.** Differentially expressed genes in CD8 and CD4 MAIT cells. The y-axis uses log FDR-adjusted p value with a significance level of FDR-adjusted p < 0.05. **G.** Dot plot displaying differential gene analysis between three subsets of MAIT cells; color intensity represents the average gene expression per cell and the dot size represents the frequency of cells expressing each gene.

In our CITE-seq dataset, we observed that 50% of the sorted MAIT cells were DN, 35% CD8^+^ and 15% of CD4^+^ defined by both the presence of CD4/CD8 transcript and surface protein expression[1] (**Supplemental Fig 2C,D**). Stratifying by CD4 and CD8 surface protein expression, we unexpectedly found that all MAIT cells had detectable *CD8A* transcript regardless of CD4 or CD8 surface expression and that surface CD8 expression correlated with *CD8B* transcript expression (**Fig 2E**). The CITE-seq technology also enables accurate annotation of DN MAIT cells that were CD4/CD8 surface barcode negative and neither expressed *CD8B* nor *CD4* transcript, in contrast to prior work that relied on surface protein or transcript alone and was potentially subject to “dropout” [29, 30]. C2 MAIT cells expressed CD4 surface protein and *CD4* transcript and were considered CD4^+^ MAIT cells (**Fig 2E**). DP MAIT cells (CD4^+^CD8^+^ surface barcodes) were not detected in our dataset (**Fig 2E**). We confirmed previous findings[1] that CD4^+^ and CD8^+^ MAIT cells have distinct transcriptional signatures; CD4^+^ MAIT cells differentially expressed co-stimulatory receptors *CTLA4*, *TNFRSF4*, the cytokine IL32 and the IL2 receptor α (*IL2RA*), while CD8^+^ MAIT cells differentially expressed a cytotoxic program including *NKG7*, *KLRG1*, *EOMES*, *GNLY*, PRF, *GZMK* and *TNF* (**Fig 2F, 3A, Supplemental Fig 3, Supplemental Table 2)**. DN MAIT cells differentially expressed activation-associated genes such as *JUN*, *FOS*, *DUSP1*, *XCL1*, *XCL2* and *IFITM3* (**Fig 2 F**). We next stratified our analyses by stimulation condition and found that CD8^+^ MAIT cells significantly upregulated transcription of *GZMA* and *PRF1* with *Mtb* lysates, while CD4^+^ MAIT cells upregulated *IFNG* (**Fig 3B, C**).

**Figure 3:**
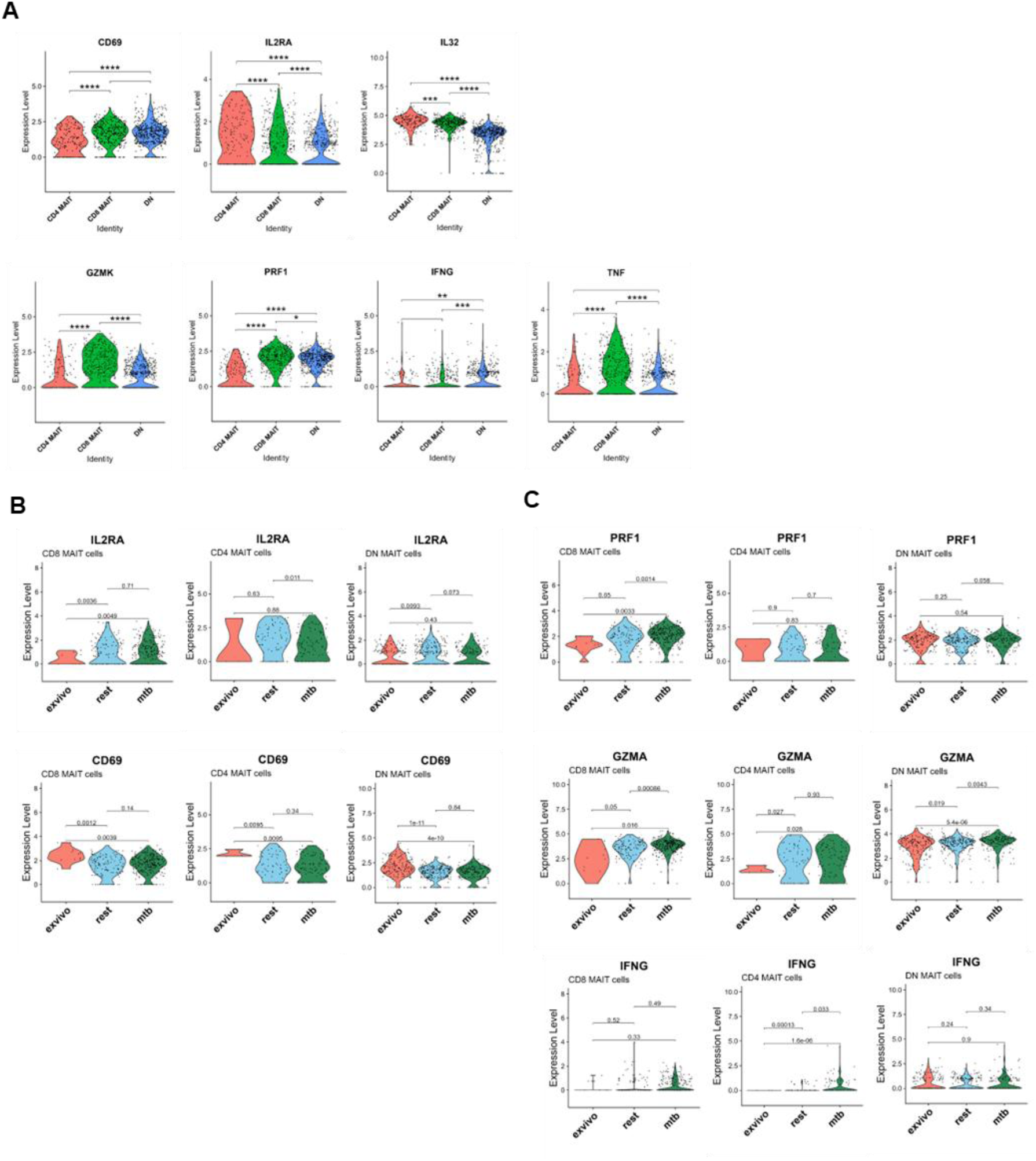
CD4+ and CD8+ MAIT cells express distinct gene signatures. Violin plots displaying gene expression levels of **A.** Activation and effector transcripts in MAIT cell subsets in pooled experimental conditions and **B, C** stratified by each in vitro stimulation condition. Statistical comparisons made by unpaired Wilcox test, p<0.05 significant, p-value adjusted.

### CD4^+^ MAIT cells express diverse TCRs

We next tested the hypothesis that these functionally distinct CD4^+^ and CD8^+^ MAIT cells adopted distinct TCR usage by integrating VDJ sequencing analyses of sorted MAIT cells from our laboratory (2 donors) with one published and publicly available dataset from *Garner et* al[25] that also employed VDJ sequencing of MR1-5-OP-RU-tetramer^+^ cells (3 donors) but restricted most of the published analysis to CD8^+^*TRAV1-2*^+^MAIT cells (**Supplemental Fig 4,5)**. MAIT cells were identified in both datasets using a combination of transcripts including *KLRB1, IL18R1, ZBTB16* and then stratified by co-expression of CD4 and CD8. Using combined datasets, we again observed that *TRAV1-2*^+^ positive MAIT cells differentially expressed cytotoxic gene signatures including *KLRB1, TRAV1-2, SLC4A10, NKG7* and *CCL3* whereas *TRAV1-2*^-^ MAIT cells differentially expressed *CD4* in addition to non-cytotoxic and co-stimulatory genes, including *ICOS, SELL, MAL, TNFRSF4* and *HLA-DRA* (**Fig 4A**).

**Figure 4:**
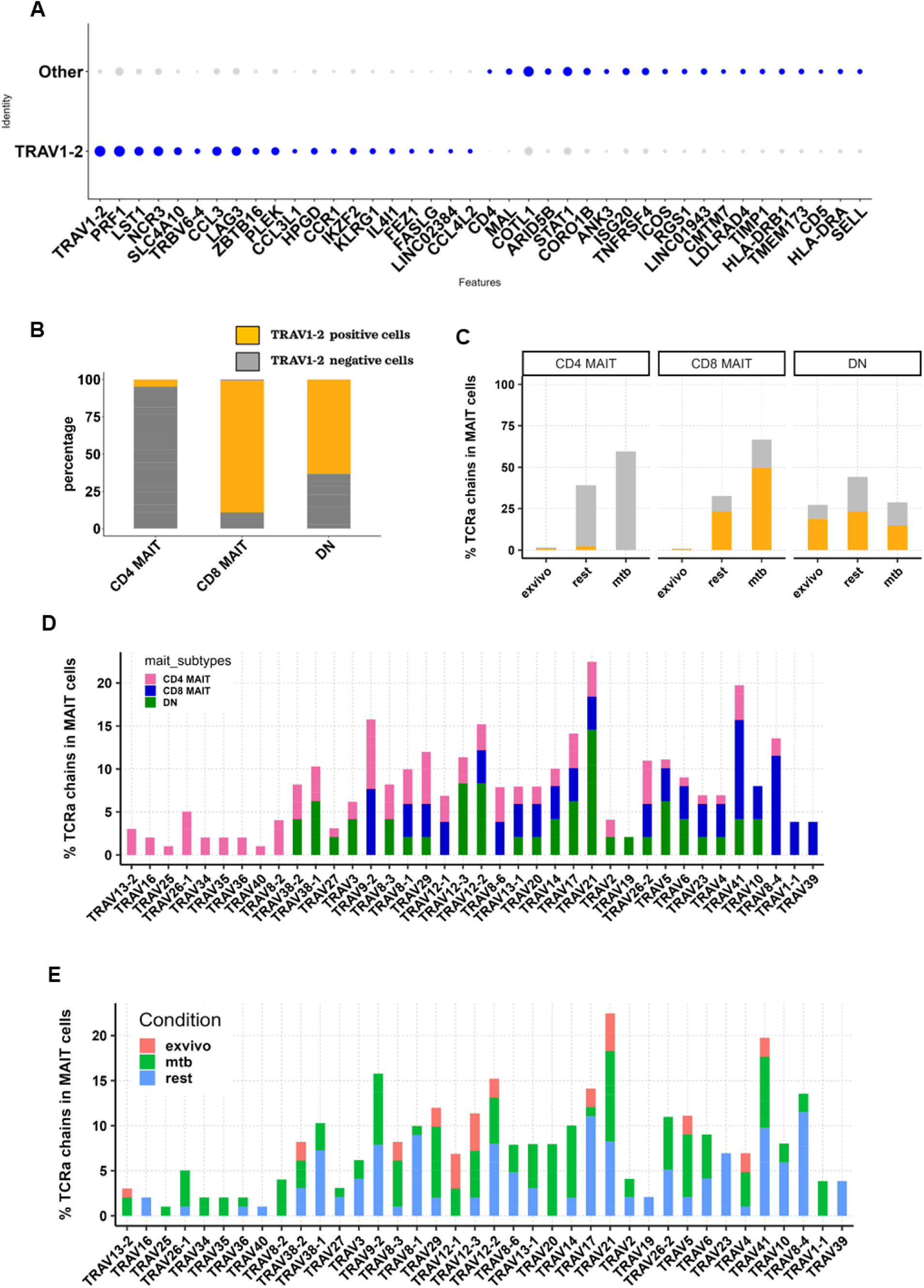
CD4+ MAIT cells predominantly express TRAV1-2 negative TCR chains. **A** Dot plot displaying the top differentially expressed genes of MR1-5-OP-RU tetramer+ TRAV1-2+ versus negative MAIT cells**. B,C.** Bar plot showing the proportion of cells expressing the canonical TRAV1-2 TCR (yellow) or TRAV1-2 negative TCRs (grey) in each MAIT cell subset. **D.** TCR*α* diversity measured in pooled experimental conditions stratified by MAIT cell subset or E. stimulation condition.

**Figure 5:**
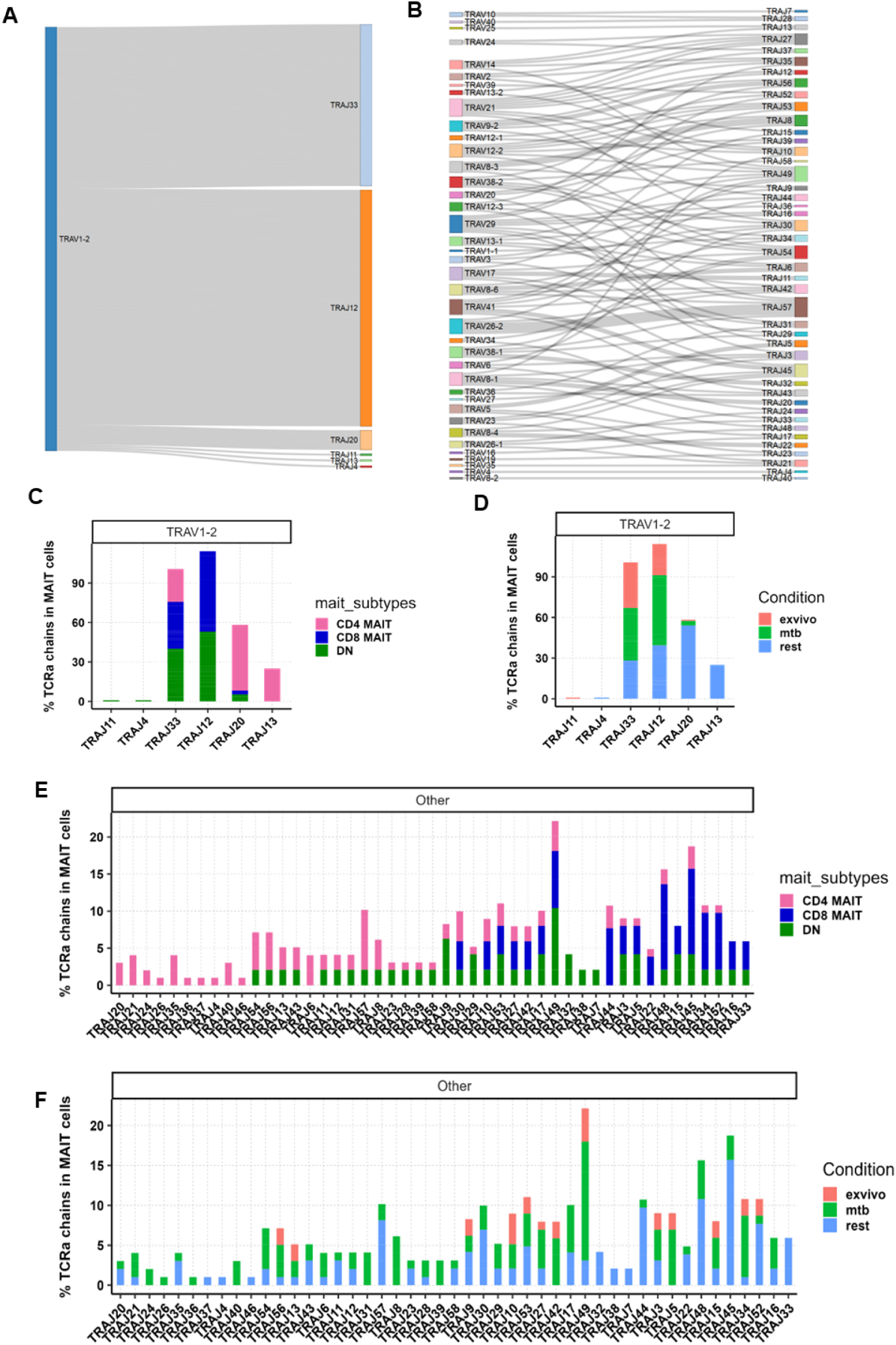
TRAJ chain diversity in MAIT cells. **A.** Sankey plot displaying TRAJ chain diversity in TRAV1-2 positive and **B.** TRAV1-2 negative MAIT cells. **C.** Bar plot displaying TRAJ chain usage stratified TRAV1-2 positive MAIT cell subset and **D.** stimulation condition. **E.** Bar plots displaying TRAJ chain usage in TRAV1-2 negative cells stratified by MAIT cell subset and **F.** stimulation condition.

We identified more than 7,587 paired TCRs in MAIT cells from five donors. First, we performed VDJ analysis of all MAIT cells irrespective of experimental condition and observed that 90% of CD4^+^ MAIT cells had non-canonical TRAV1-2^-^ TCRs relative to 70% from *Garner et al*. In contrast, only 15% of CD8^+^ and ∼35% of DN MAIT cells expressed non-canonical TRAV1-2α chains (**Fig 4B**, **6A**). TRAV1-2^-^ TCR expression increased after in vitro incubation with IL2 with or without *Mtb* lysate mainly in CD4^+^ MAIT cells (**Fig 4C**). CD4^+^ MAIT cells expressed *TRAV18-1, TRAV16, TRAV25, TRAV26-1, TRAV34, TRAV35, TRAV36, TRAV40 and TRAV8-2* (*Kaur et al*) and *TRAV29, TRAV22, TRAV30, TRAV-35 and TRAV39* (*Garner et al*). Only *TRAV35+* CD4^+^ MAIT cells were observed in both datasets. A minority of CD8^+^ MAIT cells also expressed non-canonical TRAV chains including *TRAV1-1* and *TRAV39* (*Kaur et al*) and *TRAV10* and *TRAV9-1* (*Garner et al*). The majority of DN MAIT cells expressed shared non-canonical TRAVs between CD8^+^ and CD4^+^ MAIT cells in both datasets such as *TRAV21, TRAV8-1, TRAV12-2, TRAV12-3, TRAV41, TRAV13-1, TRAV14, TRAV20, TARV17* and *TRAV5* (*Kaur et al* and *Garner et al*). In the *Kaur et al* dataset, DN MAIT cells expressed one distinct TCRα chain, TRAV*19*, not observed in CD4^+^ or CD8^+^ MAIT cells (**Fig 4D, 6B**). We also detected additional non-canonical α chains such as *TRAV8-1, TRAV21, TRAV12-2, TRAV12-3* and *TRAV4* expression in MAIT cells identified in prior studies that used targeted single cell TCR amplification[26, 31] (**Fig 4D, 6B**).

Next, we tested the hypothesis that in vitro re-stimulation could select for select TRAV1-2^-^ chain usage against tuberculosis, and we observed non-canonical TCRs in both IL2 +/- *Mtb* lysates conditions compared to ex vivo (**Fig 4E**). *Mtb* lysate induction was associated with unique TCR usage including *TRAV34, TRAV35*, *TRAV1-1, TRAV8-2,* and *TRAV25* in predominantly CD4^+^ MAIT cells (**Fig 4E**).

Additionally, we found 112 MAIT cells (9% of total) expressed dual TCRα chains (**Supplemental Fig 6**), consistent with one report where all MAIT cells expressed at least one *TRAV1-2* chain [24]. In contrast, we found 35% of dual MAIT cells expressed two isoforms of *TRAV1-2* chains with different J-segments and CDR3α sequences, 40% of MAIT cells co-expressed *TRAV1-2* with a different α chain and 25% of MAIT cells expressed two non-canonical α chains (**Supplemental Fig 6A**). The majority of CD8^+^ and DN MAIT cells expressed two *TRAV1-2* chains whereas few CD4^+^ MAIT cells with dual TCRs expressed one *TRAV1-2* chain (**Supplemental Fig 6A,B**).

Next we analyzed β chain expression in TRAV1-2^+/-^ MAIT cells and found that 90% of *TRAV1-2*^+^ MAIT cells paired with canonical β chains such as *TRBV6-4* or 6-1 or 20-1, consistent with several previous reports [32, 33]. The remaining 10% paired with non-canonical TRBV chains including *TRBV25-1, TRBV24-1, TRBV28*, *TRBV11-1* (**Supplemental Fig 7A)**. *TRAV1-2*^-^ MAIT cells demonstrated significantly more diverse TRBV usage (**Supplemental Fig 7B**), resembling private TCRs.

Next, we analyzed MAIT cell J segment diversity and observed that 90% of *TRAV1-2* positive MAIT cells had canonical J usage (*TRAJ-33/20/12*) as previously reported [32] whereas *TRAV1-2*^-^ MAIT cells demonstrated significantly more diverse J gene expression (**Fig 5A**, **B**). While TRAJ33 and 12 were the predominant J chains expressed by *TRAV1-2*^+^ MAIT cells in both datasets, we found that CD4^+^ *TRAV1-2^+^* MAIT cells largely expressed *TRAJ33* and 20 (*Kaur et al*) (**Fig 5C, D**, **6C**). Only 8% of *TRAV1-2*-MAIT cells demonstrated canonical J gene expression (TRAJ33/20/12) in both datasets consistent with previous studies [25, 32]. CD4^+^ *TRAV1-2*^-^ MAIT cells also expressed more diverse J genes relative to CD8^+^ and DN *TRAV1-2*^-^ MAIT cells (**Fig 5E).** In vitro incubation with IL2 +/- *Mtb* lysates also selected for non-canonical TRAJ usage (**Fig 5F**), similar to our observations with TRAV selection (**Fig 4E**). CD4^+^ and DN MAIT cells expressed various shared J-segments such as *TRAJ8, TRAJ 23, TRAJ28, TRAJ39, TRAJ9, TRAJ58, TRAJ57, TRAJ56, TRAJ54, TRAJ43, TRAJ11, TRAJ12, TRAJ13* and *TRAJ29* while CD8 and CD4 MAIT cells only shared two J-segment *TRAJ22* and *TRAJ44*. We observed some J-segments commonly expressed between all three MAIT cell subsets, including *TRAJ30, TRAJ10, TRAJ53, TRAJ27, TRAJ42, TRAJ49, TRAJ3, TRAJ5, TRAJ48, TRAJ45, TRAJ34* and *TRAJ52.* CD4^+^ MAIT cells expressed unique J genes including *TRAJ20, TRAJ21, TRAJ24, TRAJ26, TRAJ35, TRAJ36, TRAJ37, TRAJ4, TRAJ6, TRAJ40* and *TRAJ46* while DN TRAV1-2^-^ MAIT cells expressed more restricted chains including *TRAJ32, TRAJ38* and *TRAJ7* (*Kaur et al*). CD8^+^ MAIT cells shared non-canonical TRAJ segments with CD4^+^ and DN MAIT cells, and we did not detect any unique J segment enrichment in CD8^+^ *TRAV1-2*^-^ MAIT cells **(Fig 5E, F**). This contrasted with the *Garner et* al[25] dataset where CD4^+^ and CD8^+^ *TRAV1-2*^-^ MAIT cells demonstrated similar J chain usage, likely explained by the study’s limited sequencing and analysis of CD4^+^ MAIT cells (**Fig 6D**).

**Figure 6:**
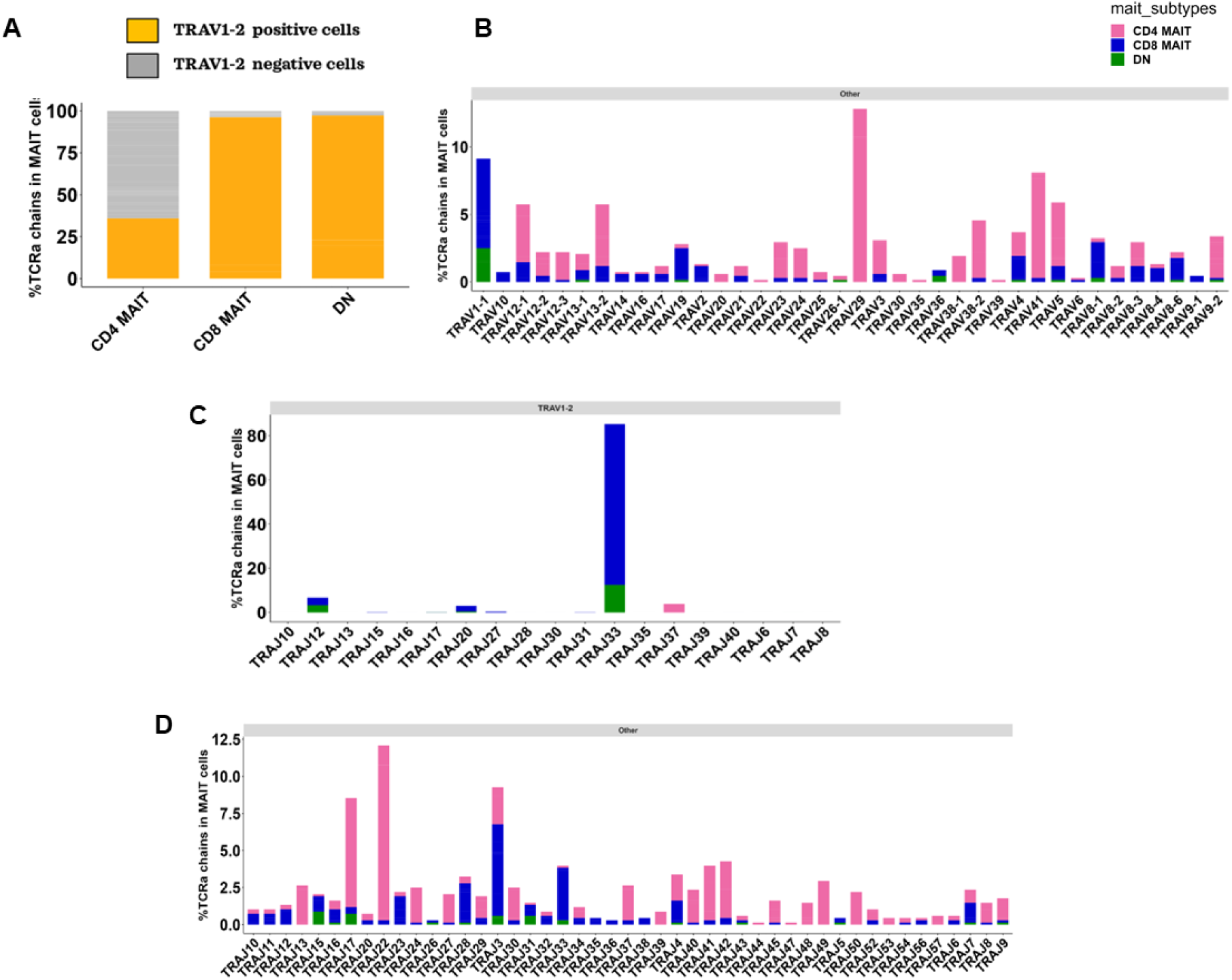
TCR diversity in MAIT cells sequenced in Garner et al. **A.** Bar plot displaying the frequency of MAIT cell subsets expressing TRAV1-2+/- TCRs. **B.** Bar plot displaying the frequency of non-canonical alpha chains expressed by MAIT cells. **C.** Bar plot displaying the frequency of J segments in canonical TRAV1-2 positive and **D.** TRAV1-2 negative MAIT cells.

### TRAV1-2 negative MAIT cells have more variable CDR3α and β sequences

Next, we compared the complementarity determine regions CDR3α and β of *TRAV1-2*^+/-^ MAIT cells that confer antigen-specificity to MR1-bound ligands. Ninety-percent of *TRAV1-2*^+^ MAIT cell CDR3α sequences were 12-14 amino acids long with a conserved tyrosine (Y) residue at the 95^th^ position in both data sets as previously reported to be essential for MR1-presented riboflavin intermediate binding[16, 34]. In contrast, CDR3α sequences of *TRAV1-2^-^* MAIT cell demonstrated more variation in the central region from positions 6-9 and about 25% of *TRAV1-2*^-^ MAIT cell subsets from both datasets expressed tri-glycine (GGG) or tetra-glycine (GGGG) motifs (**Fig 7A-E**; **Fig 8A-E**). There was also more variance in CDR3α sequence length from 7 to 23 amino acids in *TRAV1-2*^-^ MAIT cells (**Fig 7F,G; 8F-G**). Despite variable lengths, the CDR3α sequences maintained conserved terminals consisting of cysteine (C) and alanine (A) at 1 and 2 positions, respectively, with 60 to 70% containing valine (V) at the 3^rd^ position in both datasets. We found that the CDR3α sequences of all MAIT cells maintained conserved N-terminal amino acids including Leucine (L) or Isoleucine (I) or Threonine (T) or Phenylanaline (F). Next, we compared the CDR3α amino acid sequences of CD4^+^ and CD8^+^ *TRAV1-2*^-^ MAIT cells and observed that CD4^+^ *TRAV1-2^-^* MAIT cells predominantly expressed tetra-glycine (GGGG) and tetra-asparagine (NNNN) from 6^th^ to 9^th^ position, whereas TRAV1-2 negative CD8^+^ MAIT cells had tri-glycine (GGG) expression from 7^th^ to 9^th^ position (**Fig 7C-E**; **Fig 8C-E**). The CDR3α length of both CD4^+^ and CD8^+^ *TRAV1-2^-^* MAIT cells varied from 7 to 18 amino acids in our data set (*Kaur et al*) and up to 23 amino acids in *Garner et al* (**Fig. 7F, G; 8F, G)**.

**Figure 7:**
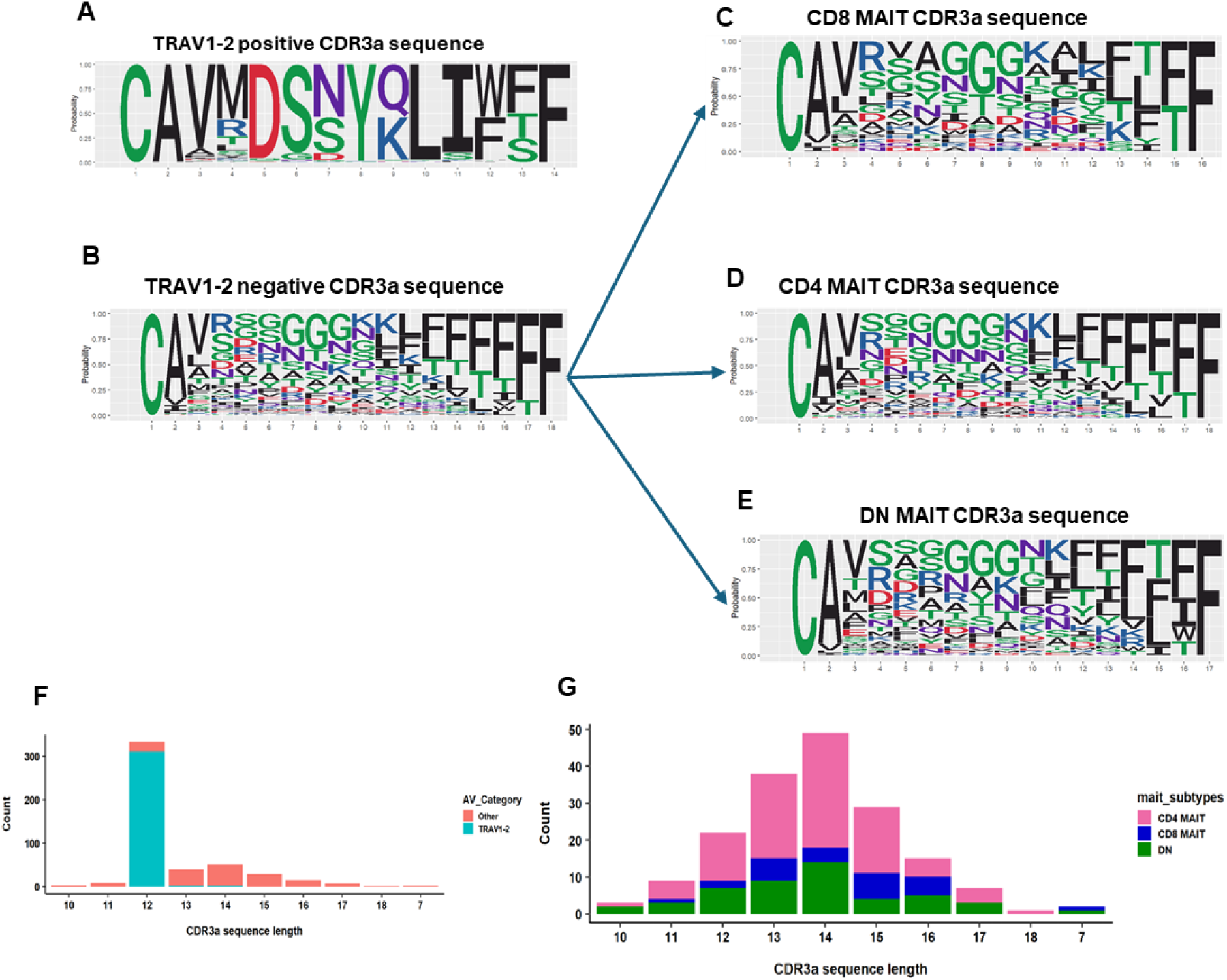
TRAV1-2 negative MAIT cell CDR3α diversity. Sequence logo plots displaying the CDR3α sequences expressed by **A.** TRAV1-2 positive MAIT cells, **B.** TRAV1-2 negative MAIT cells and **C-E.** MAIT cell subsets. **F.** Bar plot displaying the length of CDR3α sequence in TRAV1-2+/- MAIT cells. **G**. Bar plot displaying the amino acid length of CDR3α sequence in TRAV1-2 negative MAIT cells stratified by subset.

**Figure 8:**
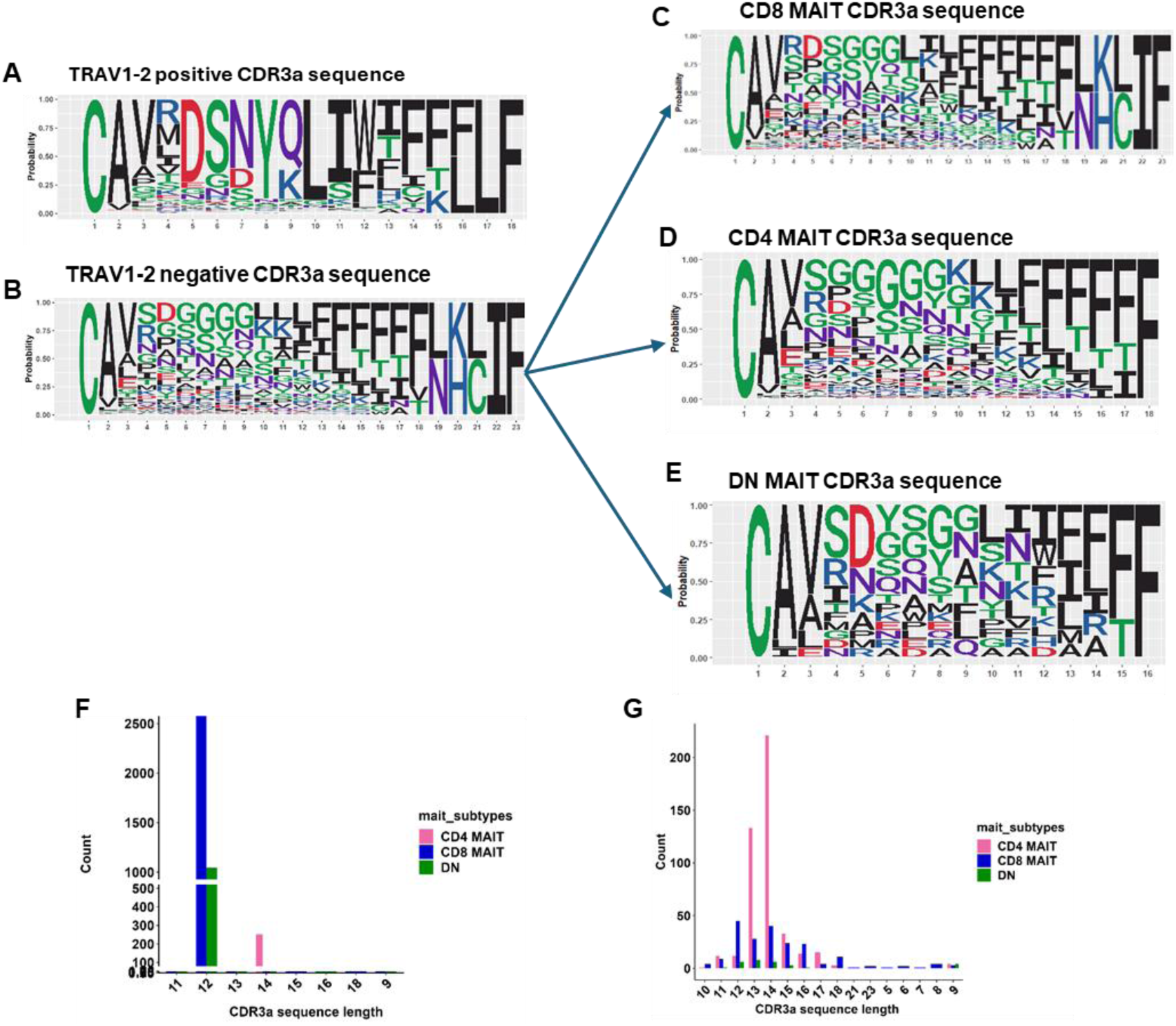
TRAV1-2 negative MAIT cell CDR3α diversity in Garner et al. Sequence logo plot displaying CDR3α sequences from Garner et al expressed by. **A.** TRAV1-2 positive MAIT cells **B.** TRAV1-2 negative MAIT cells and **C-E.** TRAV1-2 negative MAIT cell subsets **F.** Bar plot displaying the length of CDR3α sequence stratified by MAIT cell subset. **G.** Bar plot displaying the amino acid length of CDR3α sequence in TRAV1-2 negative MAIT cell subsets.

Next we compared the CDR3β sequences and did not detect significant differences between *TRAV1-2*^+/-^ MAIT cells in both datasets. The CDR3β sequence of *TRAV1-2*^+/-^ MAIT cells demonstrated conserved terminals consisting of cysteine (C) and alanine (A) at 1 and 2 positions and a string of 4 to 5 Phenylalanine (F) amino acid residues at the other end, except one CDR3β sequence with extended length in Garner et al (**Supplemental Fig 8A, B and 9A, B**). We also compared the CDR3β sequences between MAIT cell subsets and again did not detect significant differences in CD4^+^, CD8^+^ or DN MAIT cells in both datasets (**Supplemental Fig 8C-E and 9C-E**). The majority of CDR3β chains of TRAV1-2^+/-^ MAIT cells ranged between 14 to 16 amino acids (**Supplemental Fig 8F,G and 9F,G**).

### Functional annotation of MAIT cell TCRs and epitope prediction

As *TRAV1-2* negative CD4^+^ MAIT cells demonstrated unexpected TCR diversity, we hypothesized that these TCRs may bind alternative epitopes to canonical riboflavin intermediates. We performed similarity searches against publicly available database using Mc-TCR software map (http://friedmanlab.weizmann.ac.il/McPAS-TCR/) to compare *TRAV1-2^+/-^* MAIT cell CDR3α and β sequence homology with a reference library. We found that CDR3α and β sequences of CD8^+^, CD4^+^ and DN *TRAV 1-2*^+^ MAIT cells showed 100% similarity with published canonical CDR3α and β sequences associated with *Mtb* infection datasets [35, 36] while CDR3α and β sequences of CD8^+^, CD4^+^ and DN *TRAV1-2^-^* MAIT cells had only 60-80% similarity at the central region with *M. tuberculosis*-specific MAIT cells, likely due to restricted definitions of MAIT cells as TRAV1-2^+^CD8^+^ T cells in prior studies [35, 36]. These *TRAV1-2^-^*MAIT cells also demonstrated 60-80% similarity with peptide-specific epitopes derived from Cytomegalovirus (CMV) pp65 antigen and DQ2.5-glia antigen of Celiac autoimmune disease signatures (Supplemental Table 3). Modification of one amino acid in the CDR3α and β sequence in silico increased the similarity to >90% with published CDR3α and β sequences associated with *M*tb infection. Modification of two amino acids in CDR3α and β sequence further increased the similarity to 100% with published CDR3α and β sequences including peptide cancer antigens Hemoglobin-like protein (HbO), EphA2, NY-ESO-1, MART1 and T72 (Tn), Hepatitis, COVID, CMV and Epstein Barr, Influenza and Yellow fever viral antigens [35–39] (Supplemental Table 3).

We next performed Gene Ontology (GO) annotations using Kyoto Encyclopedia of Genes and Genomes (KEGG) and string networking analysis that demonstrated that *TRAV1-2*^+^ and *TRAV1-2*^-^ MAIT cells associated with different functional pathways (**Fig 9**). Our analysis revealed that TRAV1-2^+^ MAIT cells expressed signatures of cell-mediated immunity whereas *TRAV1-2*^-^ MAIT cells expressed genes involved in metabolic control and immunoglobulin production pathways.

**Figure 9:**
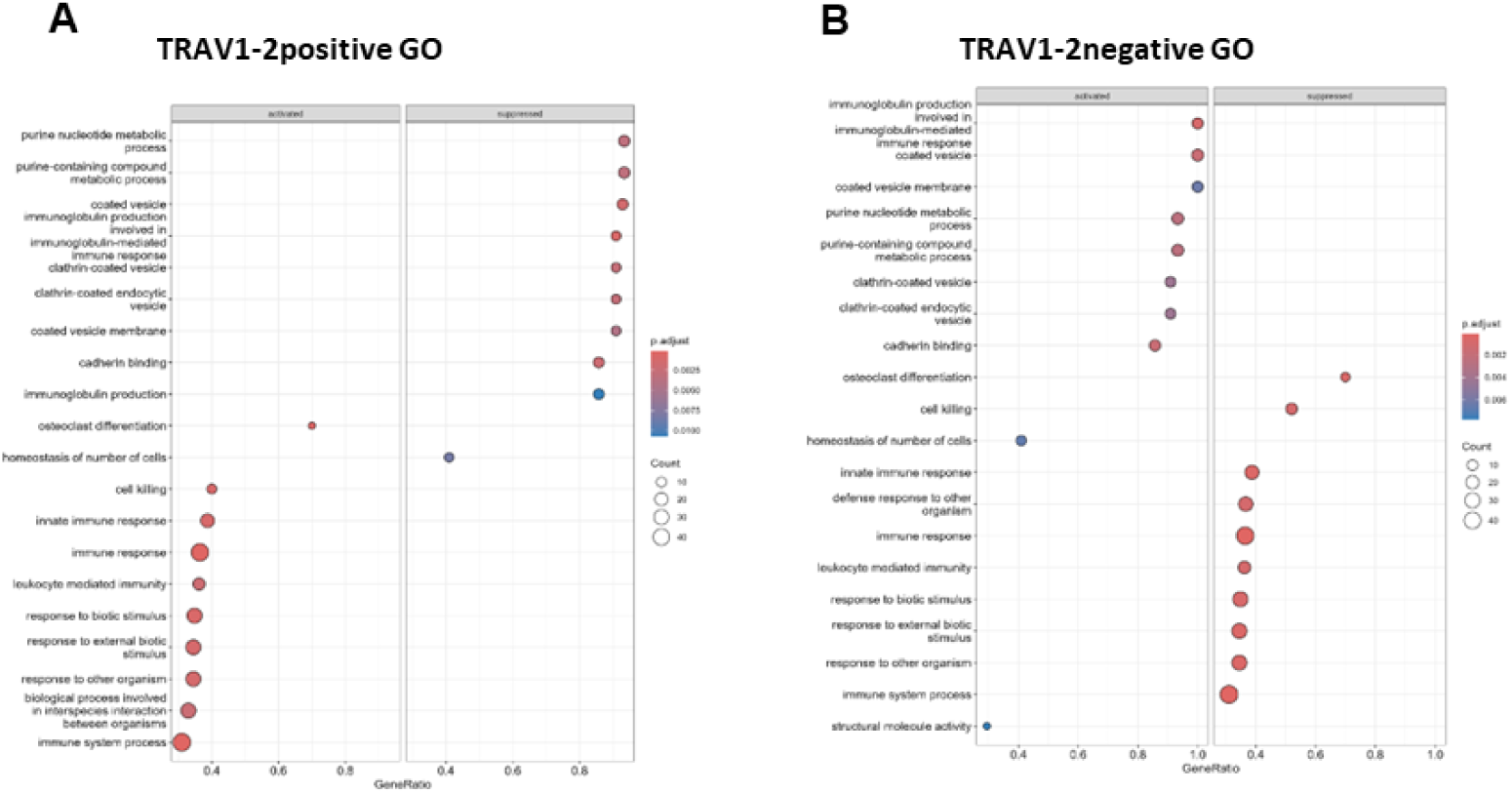
Functional annotations of TRAV1-2+/- MAIT cells. Dot plot displaying functional phenotypes of **A.** TRAV1-2 positive **B.** TRAV1-2 negative MAIT cells annotated in the Gene Ontology (GO) database using Rstudio.

## Discussion

Despite highly conserved CD8^+^TRAV1-2^+^ TCRs expressed by most 5-OP-RU-reactive MAIT cells, there is increasing appreciation of diverse TRAV1-2^-^ MAIT cells [34] with distinct functional phenotypes [1, 3, 24]. Our major finding in the present study is that the TCR co-receptor, CD4, is a major feature of these non-canonical MAIT cells in humans and that these CD4+ TRAV1-2 negative MAIT cells resemble private TCRs. Several outstanding questions remain about how CD4^+^ and other TRAV1-2^-^ MAIT cells are selected in vivo as they have lower affinity for riboflavin intermediates such as 5-OP-RU and cannot easily be studied in animal models as mature murine or non-human primate MAIT cells do not express significant levels of CD4 protein [2]. Together, these data suggest that CD4^+^ MAIT cells may be selected for in *Homo sapiens* and warrant further study in epidemiologically relevant models [1, 3, 24, 40]. While canonical TRAV1-2^+^CD8^+^ MAIT cells are well-studied and known to strongly bind and react to intermediates of riboflavin metabolism through oligoclonal J (*TRAJ33/12/20*) and TRBV chain (TRBV20/6) usage [41], the TCR repertoire of *TRAV1-2*^-^ CD4^+/-^ and CD8^+/-^ MAIT cells is not well understood. To address this gap in knowledge, we employed unbiased single-cell TCR sequencing of total human blood MAIT cells defined by MR1-5-OP-RU tetramers and stratified analyses by CD4 and CD8 surface epitope mapping coupled to transcript expression and discovered unexpected diversity in TCRα, J segment and β chain usage among non-canonical CD4^+^*TRAV1-2*^-^ MAIT cells. This work presents the first unbiased analysis of human MAIT cell TCR diversity and provides evidence to support distinct TCR usage by CD4^+^ MAIT cells suggesting alternative ligand recognition beyond 5-OP-RU or other riboflavin-derived ligands.

While CD4^+^ MAIT cells compose a minor subset of MR1-5-OP-RU tetramer^+^ cells (e,g 5-10%) [1], our finding are of high clinical significance as they are reported to be enriched during disease states including *Mtb* infection and cancer [1, 3, 24, 40] and express distinct co-stimulatory receptors including TNFRSF receptors and the master regulatory transcription factor FOXP3, with striking resemblance to conventional CD4^+^ Treg cells at both the transcript and protein level [1, 40]. Our results extend prior work by elucidating the antigen-specific binding sequences of TRAV1-2^-^ CD4^+^ MR1-5-OP-RU tetramer^+^ T cells underlying decreased reactivity to 5-OP-RU relative to TRAV1-2^+^CD8^+^ MAIT cells [16, 42]. While one study reported 6FP-MR1 tetramer reactive TRAV1-2^-^ T cells with distinct binding geometry in the MR1 pocket compared to TRAV1-2^+^ MAIT cells [16], we did not detect a convincing folate-reactive MR1-restricted T cell population nor did we observe significant expansion of TRAV1-2^+/-^ MAIT cells with folate-derived MR1 ligands. While we have observed the presence of MR1-6FP tetramer-reactive T cells by flow cytometry, they are of low and variable prevalence across human donors and do not expand using folate-derived MR1 ligand priming in vitro.

Our study provides important insights into the TCR repertoire of human CD4^+^ and CD8^+^ MAIT cells that raises critical questions as to how TRAV1-2^-^ CD4^+^ MAIT cells may recognize biochemically distinct ligands. Despite differences in immune phenotypes of CD4^+^ MAIT cells including constitutive expression of *IL2RA* (CD25), *MAL, CTLA4,* and *TNFRSF4* relative to CD8^+^ and DN MAIT cells [1], they likely do not represent a distinct developmental lineage as defined by conventional T cell ontogeny (eg, *ThPOK, RUNX3*) and may instead undergo post-thymic selection [1, 23]. A recent study in colorectal cancer found tumor-infiltrating MAIT cells enriched with CD4^+^FOXP3^+^CD39^+^ MAIT cells [40] which along with our studies in *Mycobacterium tuberculosis* [1] suggest specialized functions for CD4^+^ MAIT cells beyond cytotoxicity that may represent tissue-specific responses.

Prior structural modelling studies of non-canonical MAIT cell TCRs suggested the alternative binding affinities and geometries of TCRs with MR1 receptor such as TRAV12-2/TRBV29-1 on MAIT cells have lower binding affinity for MR1-5-OP-RU tetramers compared to cells expressing a canonical TCR (TRAV1-2/TRBV6-1) [[14, 41]. The TRAV1-2^+^ MAIT cell TCR binds to the A′ pocket of the MR1 receptor whereas TRAV1-2^-^ MAIT TCRs adopt alternative binding positions more towards the F’ pocket on MR1 [34]. These alternative binding models of non-canonical TCRs highlight challenges to identifying and selectively expanding CD4^+^ MAIT cells with diverse TCRs since they have lower affinity for the riboflavin antigens typically used to stimulate MAIT cells in vitro. A separate study found MAIT cells with higher binding affinity for MR1-5-OP-RU tetramer predominantly expressed canonical TRAV1-2 TCRs, while those with lower affinity exhibited non-canonical TCRs [26].

Importantly, there is data to support that CDR3α variability in MAIT cells can confer pathogen specificity, despite highly conserved α chains. A recent study reported a significant expansion of TRAV1-2 negative, but not TRAV1-2^+^, MAIT cells in high-risk contacts of TB cases who appear to resist primary infection, but this study did not interrogate TRAV1-2 negative chain diversity nor its association with CD4 [24]. In the present study, we discovered that the antigen-specific CDR3α sequences of TRAV1-2 negative MAIT cells were highly variable with tri or tetra Glycine (G) motifs. Together with one previous study demonstrating variable expression of glycine residues within the CDR3α region of MAIT cells conferring differential antigen recognition to *Candida* species[7], our findings further support that non-canonical TCRs may confer broader antigen-specificity beyond riboflavin metabolites.

We acknowledge a small sample size of donors as a limitation to our study, but this is mitigated by high input sorted MAIT cell sequencing across two distinct datasets that support our conclusions. Future studies of human blood and tissue-specific MAIT cells across diverse human populations in health and disease states will further inform the mechanisms by which *TRAV1-2*^-^ CD4^+^ MAIT cells are selected.

## Author contributions

The following authors contributed to the manuscript in various ways. Each author’s contribution is listed below.

Conceptualization: RK, CKV

Methodology: RK, NM, AP, DX, JA, KL, BR, DC, CKV

Investigation: RK, NM, AP, CKV

Visualization: RK, NM, AP, CKV

Funding acquisition: CKV

Writing – original draft: RK, CKV

Writing – review & editing: all authors

## Acknowledgments

We are thankful for the support of the Stony Brook Foundation, the Department of Medicine of the Renaissance School of Medicine and the Office of the Vice President of Research of Stony Brook University.

## Data and materials availability

All data, code, and materials used in the analysis will be available to any researcher for purposes of reproducing or extending the analysis.

## References

1. Vorkas CK, Krishna C, Li K, Aube J, Fitzgerald DW, Mazutis L, Leslie CS, Glickman MS: Single-Cell Transcriptional Profiling Reveals Signatures of Helper, Effector, and Regulatory MAIT Cells during Homeostasis and Activation. J Immunol 2022, 208(5):1042–1056.

2. Vorkas CK, Levy O, Skular M, Li K, Aube J, Glickman MS: Efficient 5-OP-RU-Induced Enrichment of Mucosa-Associated Invariant T Cells in the Murine Lung Does Not Enhance Control of Aerosol Mycobacterium tuberculosis Infection. Infect Immun 2020, 89(1).

3. Vorkas CK, Wipperman MF, Li K, Bean J, Bhattarai SK, Adamow M, Wong P, Aube J, Juste MAJ, Bucci V et al: Mucosal-associated invariant and gammadelta T cell subsets respond to initial Mycobacterium tuberculosis infection. JCI Insight 2018, 3(19).

4. Godfrey DI, Koay HF, McCluskey J, Gherardin NA: The biology and functional importance of MAIT cells. Nat Immunol 2019, 20:1110–1128.

5. Chen Z, Wang H, D’Souza C, Sun S, Kostenko L, Eckle SB, Meehan BS, Jackson DC, Strugnell RA, Cao H et al: Mucosal-associated invariant T-cell activation and accumulation after in vivo infection depends on microbial riboflavin synthesis and co-stimulatory signals. Mucosal Immunol 2017, 10(1):58–68.

6. Gold MC, Cerri S, Smyk-Pearson S, Cansler ME, Vogt TM, Delepine J, Winata E, Swarbrick GM, Chua WJ, Yu YY et al: Human mucosal associated invariant T cells detect bacterially infected cells. PLoS Biol 2010, 8(6):e1000407.

7. Gold MC, McLaren JE, Reistetter JA, Smyk-Pearson S, Ladell K, Swarbrick GM, Yu YY, Hansen TH, Lund O, Nielsen M et al: MR1-restricted MAIT cells display ligand discrimination and pathogen selectivity through distinct T cell receptor usage. J Exp Med 2014, 211(8):1601–1610.

8. Keller AN, Corbett AJ, Wubben JM, McCluskey J, Rossjohn J: MAIT cells and MR1-antigen recognition. Curr Opin Immunol 2017, 46:66–74.

9. Kjer-Nielsen L, Patel O, Corbett AJ, Le Nours J, Meehan B, Liu L, Bhati M, Chen Z, Kostenko L, Reantragoon R et al: MR1 presents microbial vitamin B metabolites to MAIT cells. Nature 2012, 491(7426):717–723.

10. Seshadri C, Thuong NT, Mai NT, Bang ND, Chau TT, Lewinsohn DM, Thwaites GE, Dunstan SJ, Hawn TR: A polymorphism in human MR1 is associated with mRNA expression and susceptibility to tuberculosis. Genes Immun 2017, 18(1):8–14.

11. Li K, Vorkas CK, Chaudhry A, Bell DL, Willis RA, Rudensky A, Altman JD, Glickman MS, Aube J: Synthesis, stabilization, and characterization of the MR1 ligand precursor 5-amino-6-D-ribitylaminouracil (5-A-RU). PLoS One 2018, 13(2):e0191837.

12. Rahimpour A, Koay HF, Enders A, Clanchy R, Eckle SB, Meehan B, Chen Z, Whittle B, Liu L, Fairlie DP et al: Identification of phenotypically and functionally heterogeneous mouse mucosal-associated invariant T cells using MR1 tetramers. J Exp Med 2015, 212(7):1095–1108.

13. Chancellor A, Alan Simmons R, Khanolkar RC, Nosi V, Beshirova A, Berloffa G, Colombo R, Karuppiah V, Pentier JM, Tubb V et al: Promiscuous recognition of MR1 drives self-reactive mucosal-associated invariant T cell responses. J Exp Med 2023, 220(9).

14. Gherardin NA, Souter MN, Koay HF, Mangas KM, Seemann T, Stinear TP, Eckle SB, Berzins SP, d’Udekem Y, Konstantinov IE et al: Human blood MAIT cell subsets defined using MR1 tetramers. Immunol Cell Biol 2018, 96(5):507–525.

15. Suliman S, Gela A, Mendelsohn SC, Iwany SK, Tamara KL, Mabwe S, Bilek N, Darboe F, Fisher M, Corbett AJ et al: Peripheral Blood Mucosal-Associated Invariant T Cells in Tuberculosis Patients and Healthy Mycobacterium tuberculosis-Exposed Controls. J Infect Dis 2020, 222(6):995–1007.

16. Gherardin NA, Keller AN, Woolley RE, Le Nours J, Ritchie DS, Neeson PJ, Birkinshaw RW, Eckle SBG, Waddington JN, Liu L et al: Diversity of T Cells Restricted by the MHC Class I-Related Molecule MR1 Facilitates Differential Antigen Recognition. Immunity 2016, 44(1):32–45.

17. Mak JY, Xu W, Reid RC, Corbett AJ, Meehan BS, Wang H, Chen Z, Rossjohn J, McCluskey J, Liu L et al: Stabilizing short-lived Schiff base derivatives of 5-aminouracils that activate mucosal-associated invariant T cells. Nat Commun 2017, 8:14599.

18. Patel O, Kjer-Nielsen L, Le Nours J, Eckle SB, Birkinshaw R, Beddoe T, Corbett AJ, Liu L, Miles JJ, Meehan B et al: Recognition of vitamin B metabolites by mucosal-associated invariant T cells. Nat Commun 2013, 4:2142.

19. Greene JM, Dash P, Roy S, McMurtrey C, Awad W, Reed JS, Hammond KB, Abdulhaqq S, Wu HL, Burwitz BJ et al: MR1-restricted mucosal-associated invariant T (MAIT) cells respond to mycobacterial vaccination and infection in nonhuman primates. Mucosal Immunol 2017, 10(3):802–813.

20. Kauffman KD, Sallin MA, Hoft SG, Sakai S, Moore R, Wilder-Kofie T, Moore IN, Sette A, Arlehamn CSL, Barber DL: Limited Pulmonary Mucosal-Associated Invariant T Cell Accumulation and Activation during Mycobacterium tuberculosis Infection in Rhesus Macaques. Infect Immun 2018, 86(12).

21. Souter MNT, Awad W, Li S, Pediongco TJ, Meehan BS, Meehan LJ, Tian Z, Zhao Z, Wang H, Nelson A et al: CD8 coreceptor engagement of MR1 enhances antigen responsiveness by human MAIT and other MR1-reactive T cells. J Exp Med 2022, 219(9).

22. Kurioka A, Jahun AS, Hannaway RF, Walker LJ, Fergusson JR, Sverremark-Ekstrom E, Corbett AJ, Ussher JE, Willberg CB, Klenerman P: Shared and Distinct Phenotypes and Functions of Human CD161++ Valpha7.2+ T Cell Subsets. Front Immunol 2017, 8:1031.

23. Dias J, Leeansyah E, Sandberg JK: Multiple layers of heterogeneity and subset diversity in human MAIT cell responses to distinct microorganisms and to innate cytokines. Proc Natl Acad Sci U S A 2017, 114(27):E5434–E5443.

24. Cross DL, Layton ED, Yu KK, Smith MT, Aguilar MS, Li S, Wilcox EC, Chapuis AG, Mayanja-Kizza H, Stein CM et al: MR1-restricted T cell clonotypes are associated with “resistance” to Mycobacterium tuberculosis infection. JCI Insight 2024, 9(9).

25. Garner LC, Amini A, FitzPatrick MEB, Lett MJ, Hess GF, Filipowicz Sinnreich M, Provine NM, Klenerman P: Single-cell analysis of human MAIT cell transcriptional, functional and clonal diversity. Nat Immunol 2023.

26. Suliman S, Kjer-Nielsen L, Iwany SK, Lopez Tamara K, Loh L, Grzelak L, Kedzierska K, Ocampo TA, Corbett AJ, McCluskey J et al: Dual TCR-alpha Expression on Mucosal-Associated Invariant T Cells as a Potential Confounder of TCR Interpretation. J Immunol 2022, 208(6):1389–1395.

27. Tickotsky N, Sagiv T, Prilusky J, Shifrut E, Friedman N: McPAS-TCR: a manually curated catalogue of pathology-associated T cell receptor sequences. Bioinformatics 2017, 33(18):2924–2929.

28. Shi J, Zhou J, Zhang X, Hu W, Zhao JF, Wang S, Wang FS, Zhang JY: Single-Cell Transcriptomic Profiling of MAIT Cells in Patients With COVID-19. Front Immunol 2021, 12:700152.

29. Qiu P: Embracing the dropouts in single-cell RNA-seq analysis. Nat Commun 2020, 11(1):1169.

30. Dias J, Boulouis C, Gorin JB, van den Biggelaar R, Lal KG, Gibbs A, Loh L, Gulam MY, Sia WR, Bari S et al: The CD4(-)CD8(-) MAIT cell subpopulation is a functionally distinct subset developmentally related to the main CD8(+) MAIT cell pool. Proc Natl Acad Sci U S A 2018, 115(49):E11513–E11522.

31. van Schaik B, Klarenbeek P, Doorenspleet M, van Kampen A, Moody DB, de Vries N, Van Rhijn I: Discovery of invariant T cells by next-generation sequencing of the human TCR alpha-chain repertoire. J Immunol 2014, 193(10):5338–5344.

32. Held K, Beltran E, Moser M, Hohlfeld R, Dornmair K: T-cell receptor repertoire of human peripheral CD161hiTRAV1-2+ MAIT cells revealed by next generation sequencing and single cell analysis. Hum Immunol 2015, 76(9):607–614.

33. Reantragoon R, Corbett AJ, Sakala IG, Gherardin NA, Furness JB, Chen Z, Eckle SB, Uldrich AP, Birkinshaw RW, Patel O et al: Antigen-loaded MR1 tetramers define T cell receptor heterogeneity in mucosal-associated invariant T cells. J Exp Med 2013, 210(11):2305–2320.

34. Awad W, Meermeier EW, Sandoval-Romero ML, Le Nours J, Worley AH, Null MD, Liu L, McCluskey J, Fairlie DP, Lewinsohn DM et al: Atypical TRAV1-2(-) T cell receptor recognition of the antigen-presenting molecule MR1. J Biol Chem 2020, 295(42):14445–14457.

35. Huang H, Sikora MJ, Islam S, Chowdhury RR, Chien YH, Scriba TJ, Davis MM, Steinmetz LM: Select sequencing of clonally expanded CD8(+) T cells reveals limits to clonal expansion. Proc Natl Acad Sci U S A 2019, 116(18):8995–9001.

36. Huang H, Wang C, Rubelt F, Scriba TJ, Davis MM: Analyzing the Mycobacterium tuberculosis immune response by T-cell receptor clustering with GLIPH2 and genome-wide antigen screening. Nat Biotechnol 2020, 38(10):1194–1202.

37. Glanville J, Huang H, Nau A, Hatton O, Wagar LE, Rubelt F, Ji X, Han A, Krams SM, Pettus C et al: Identifying specificity groups in the T cell receptor repertoire. Nature 2017, 547(7661):94–98.

38. Huth A, Liang X, Krebs S, Blum H, Moosmann A: Antigen-Specific TCR Signatures of Cytomegalovirus Infection. J Immunol 2019, 202(3):979–990.

39. Lee ES, Thomas PG, Mold JE, Yates AJ: Identifying T Cell Receptors from High-Throughput Sequencing: Dealing with Promiscuity in TCRalpha and TCRbeta Pairing. PLoS Comput Biol 2017, 13(1):e1005313.

40. Li S, Simoni Y, Becht E, Loh CY, Li N, Lachance D, Koo SL, Lim TP, Tan EKW, Mathew R et al: Human Tumor-Infiltrating MAIT Cells Display Hallmarks of Bacterial Antigen Recognition in Colorectal Cancer. Cell Rep Med 2020, 1(3):100039.

41. Wong EB, Gold MC, Meermeier EW, Xulu BZ, Khuzwayo S, Sullivan ZA, Mahyari E, Rogers Z, Kloverpris H, Sharma PK et al: TRAV1-2(+) CD8(+) T-cells including oligoconal expansions of MAIT cells are enriched in the airways in human tuberculosis. Commun Biol 2019, 2:203.

42. Meermeier EW, Laugel BF, Sewell AK, Corbett AJ, Rossjohn J, McCluskey J, Harriff MJ, Franks T, Gold MC, Lewinsohn DM: Human TRAV1-2-negative MR1-restricted T cells detect S. pyogenes and alternatives to MAIT riboflavin-based antigens. Nat Commun 2016, 7:12506.

