## Supplemental Figures for "CD4^+^ Mucosal-associated Invariant T (MAIT) cells express highly diverse T cell receptors"

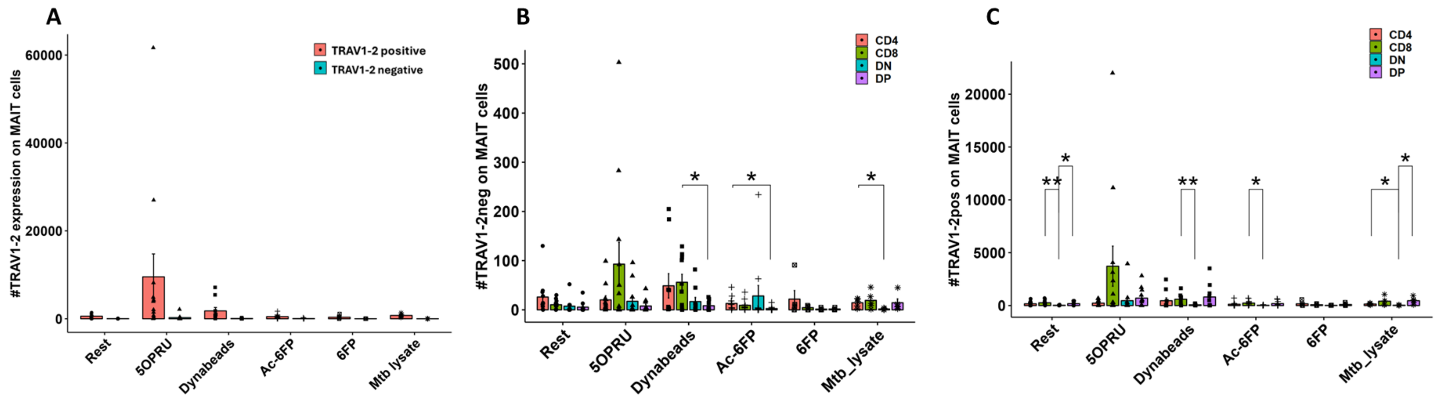


**Supplemental Figure 1: TRAV1-2 negative TCRs are present across conditions with modest expansion with 5-OP-RU stimulation. A.** MAIT cell absolute numbers stratified by TRAV1-2 expression after 7 days of incubation in vitro with various stimuli. **B.** TRAV1-2 negative (left)and positive (right) MAIT cell absolute numbers stratified by subset. Statistical comparisons made by unpaired t-test. *p<0.05

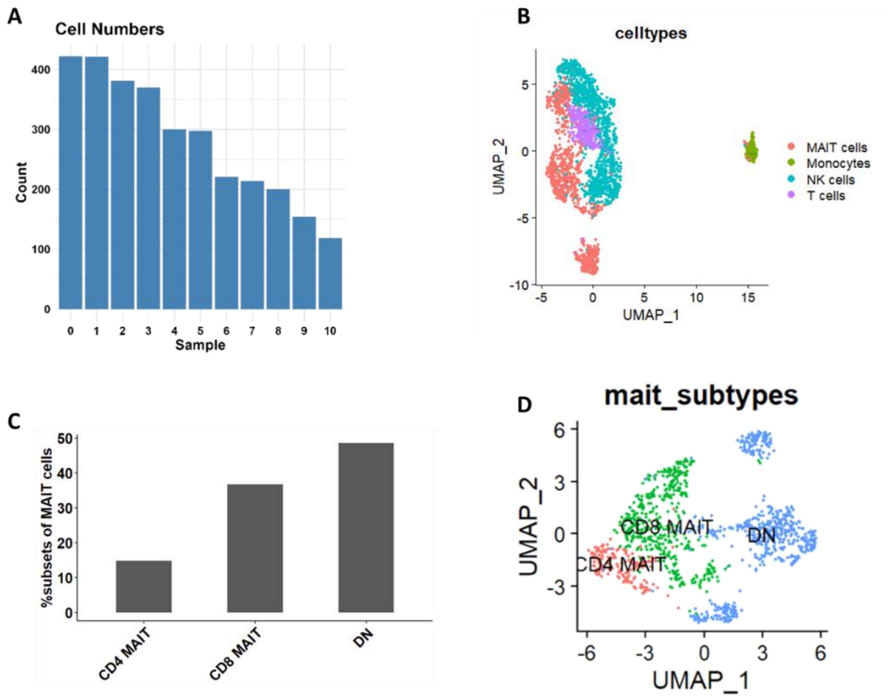


**Supplemental Figure 2: Quality control and analysis of CITE-seq data. A.** Cell count in each cluster. **B.** UMAP visualization of four cell populations. **C.** The frequency of MAIT cell subsets **D.** UMAP visualization of MAIT cell subsets.

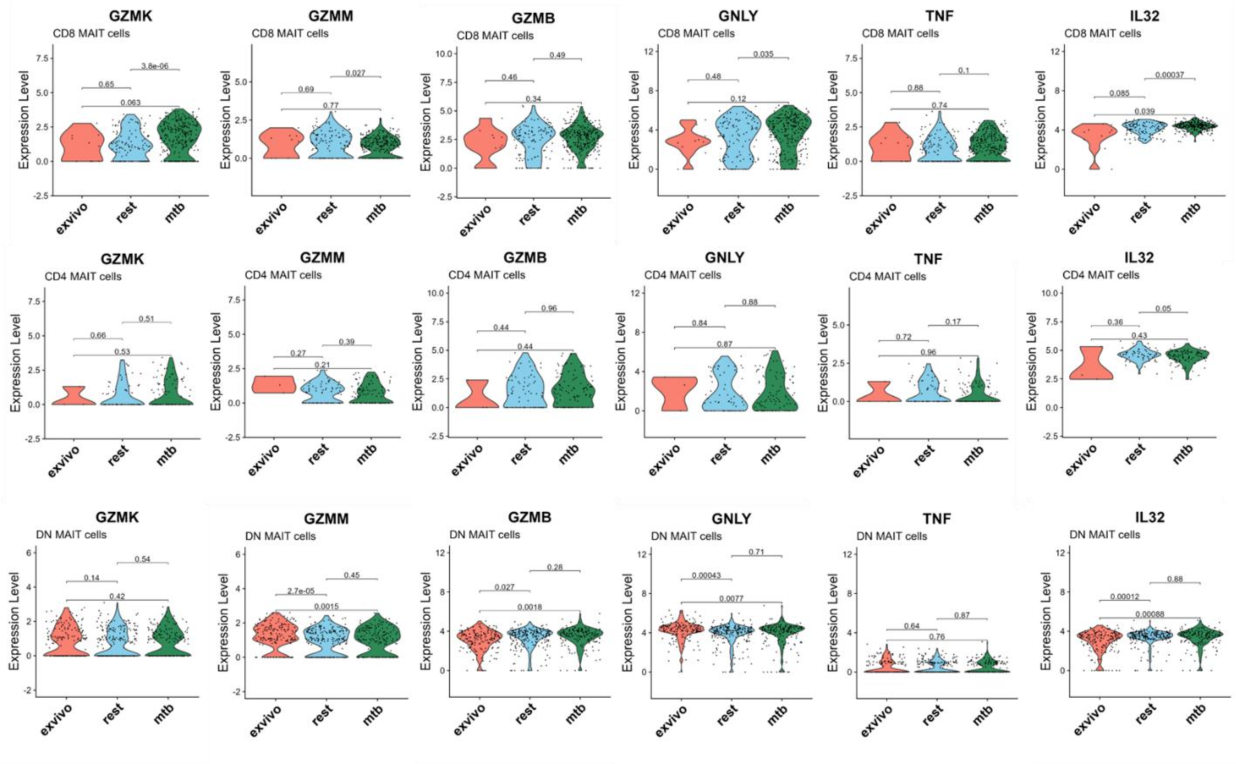


**Supplemental Figure 3: Select gene expression analysis after *Mtb* lysate induction.** Violin plots displaying the expression levels of select genes in MAIT cell subsets after co-incubation for 7 days with *Mtb* lysates relative to controls. Statistical comparisons made by unpaired Wilcox test with reported adjusted p-values and significance level of $p<0.05$.

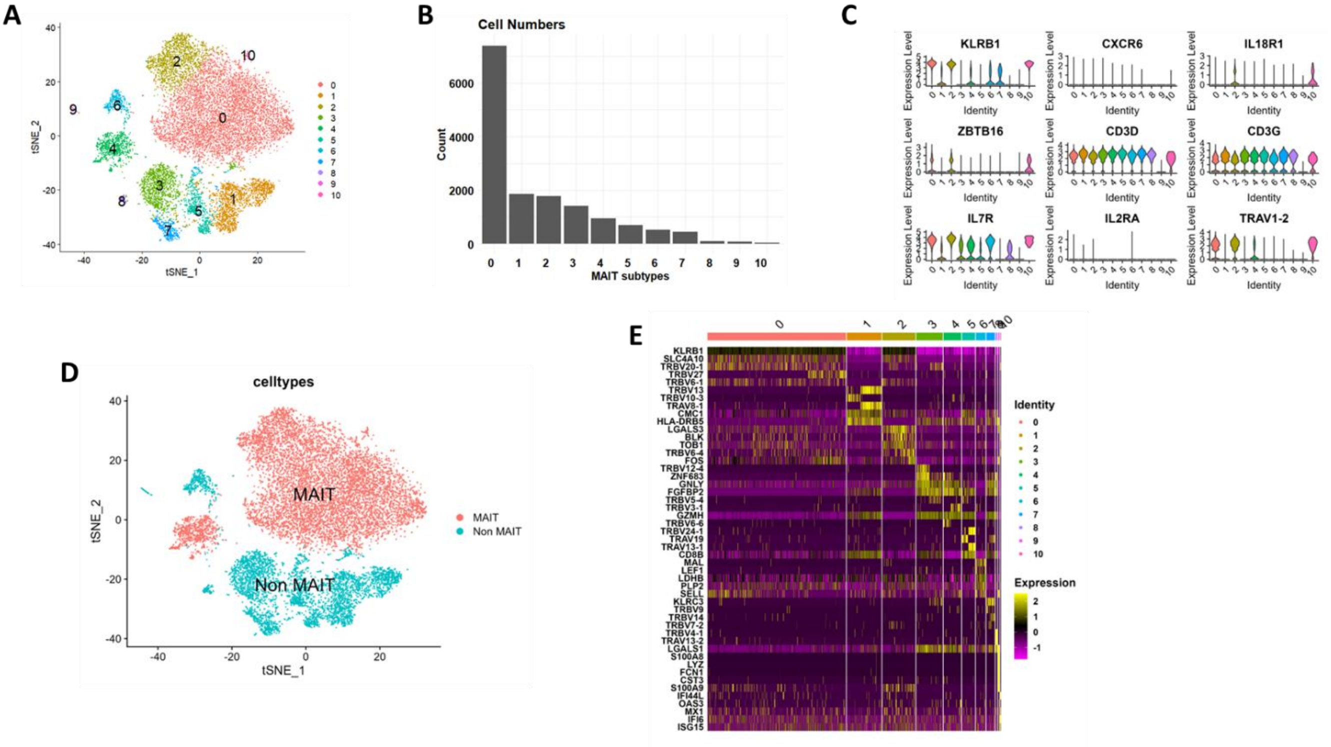


**Supplemental Figure 4: Visualization and gene expression profiling of cells from Garner et al., 2023**. **A** TSNE visualization of cells. **B.** Cell count in each cluster. **C.** MAIT cell specific gene expression in different clusters. **D.** TSNE visualization to identify MAIT cells **E.** Heat map displaying differential gene expression between MAIT cell clusters.

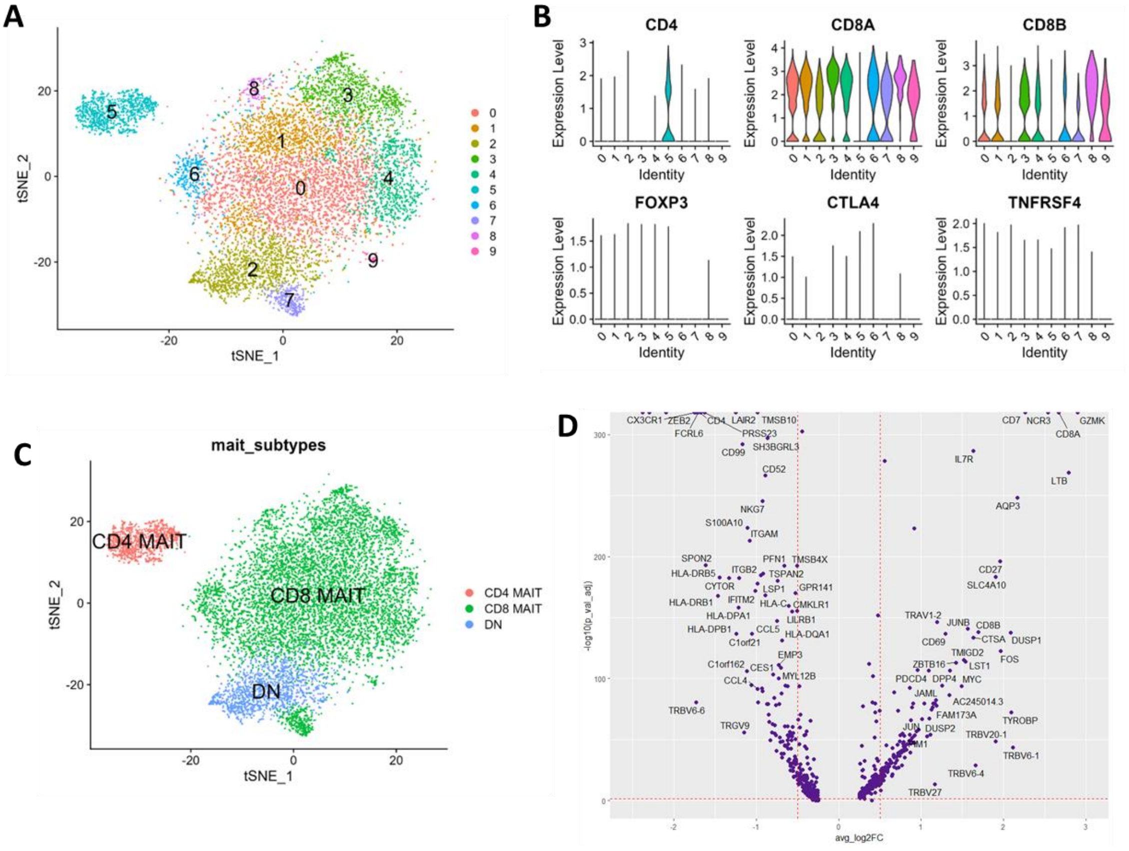

**Supplemental Figure 5: Visualization and gene expression profiling of MAIT cells (Garner et al., 2023)**. **A** TSNE visualization of MAIT cells. **B.** CD8 and CD4 co-receptor gene expression in MAIT cell clusters. **C.** TSNE visualization of MAIT subsets **D.** Volcano plot displaying differential gene expression between CD4 (-log2FC) and CD8 (+log2FC) MAIT cells.

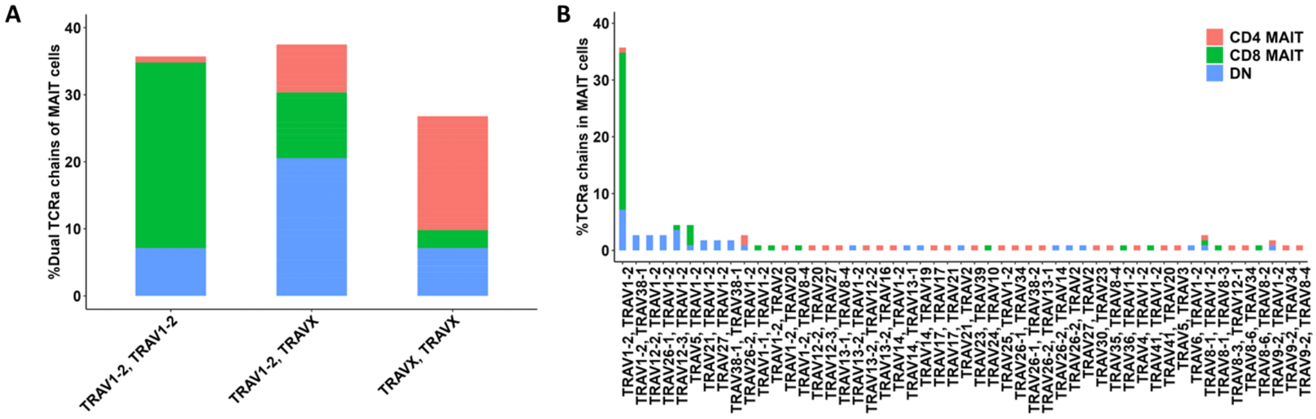


**Supplemental Figure 6: Dual TCR expression across MAIT subsets A.** Bar plot displaying combination of α chains in MAIT cells stratified by subset. TRAVX represents non-TRAV1-2 α chains. **B.** Frequency of dual TCR alpha chain combinations stratified by MAIT cell subset. Color legend at top right applies to both panels.

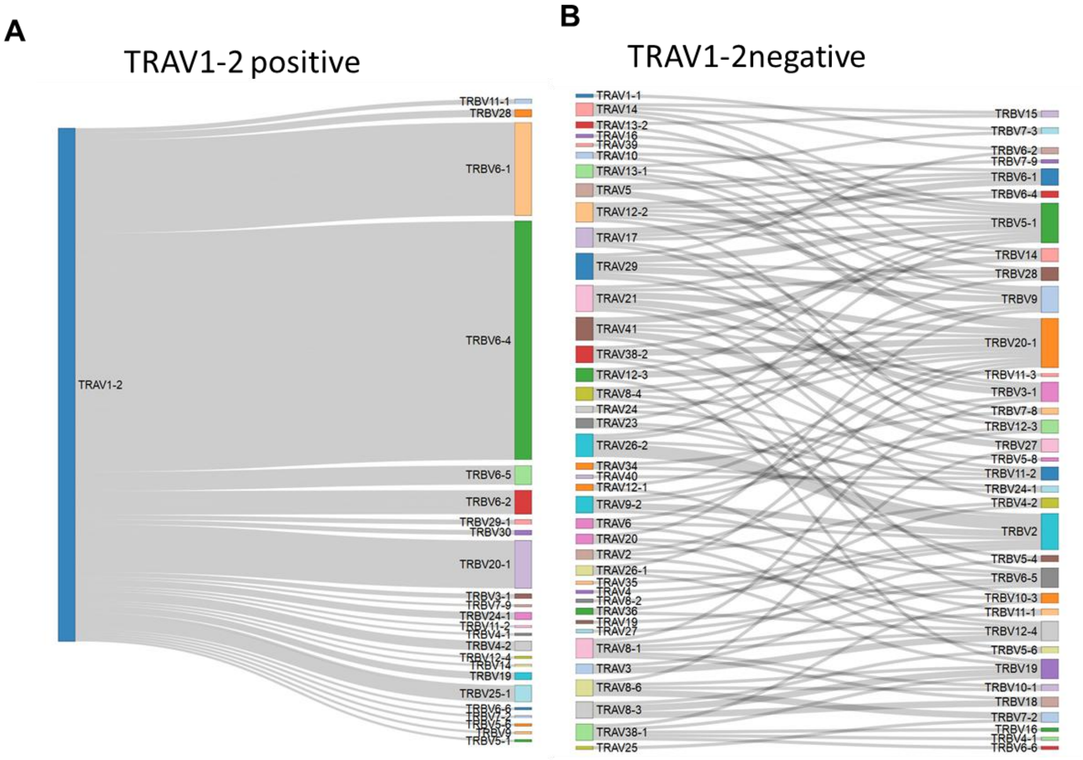


**Supplemental Figure 7: Sankey plots displaying pairing of TRAV1-2 positive (left) and TRAV1-2 negative (right) MAIT cell TCRs with TRBV chains.**

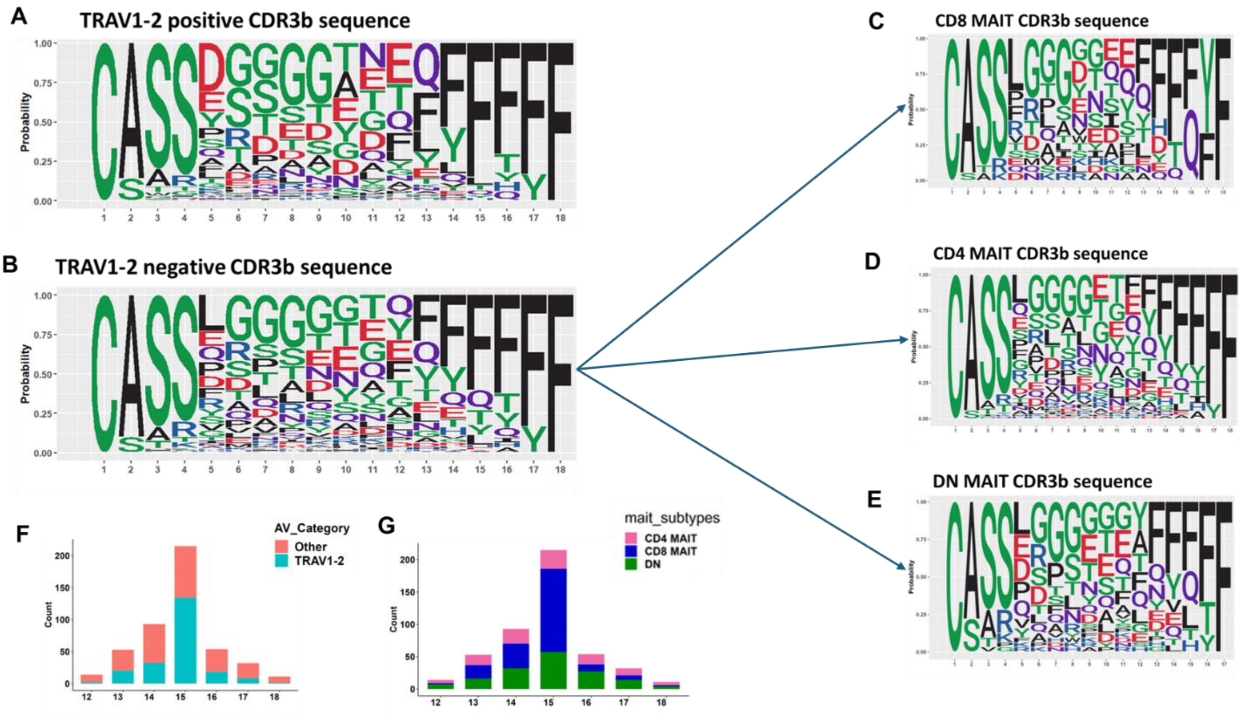


**Supplemental Figure 8: TRAV1-2+/- CDR3β sequence diversity.** Sequence logo plots displaying the CDR3β sequences expressed by **A.** *TRAV1-2*+ MAIT cells, **B.** *TRAV1-2*- MAIT cells and **C-E** MAIT cell subsets **F, G.** Bar plot displaying the amino acid length of CDR3β sequence stratified by TRAV1-2 usage **(F)** and stratified by MAIT cell subset **(G)**.

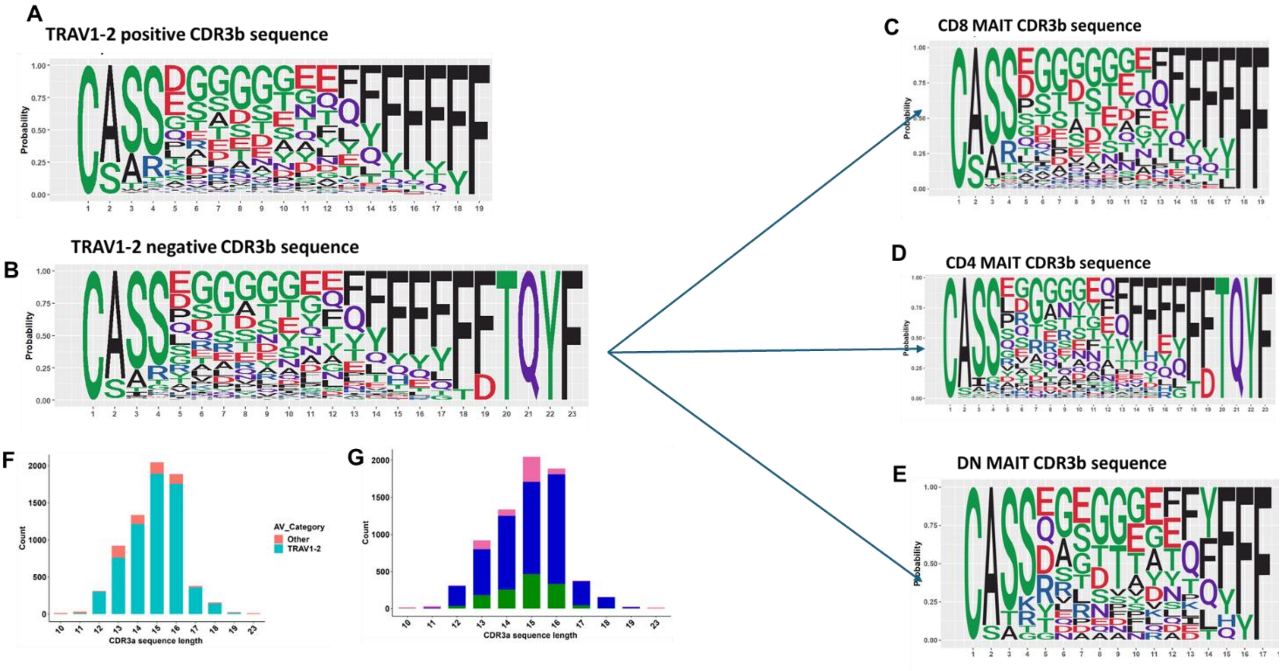


**Supplemental Figure 9: TRAV1-2+/- CDR3β sequence diversity in Garner et al.** Sequence logo plots displaying the CDR3β sequences expressed by **A.** *TRAV1-2*+ MAIT cells, **B.** *TRAV1-2*- MAIT cells and **C-E** MAIT cell subsets **F, G.** Bar plot displaying the amino acid length of CDR3β sequence stratified by TRAV1-2 usage **(F)** and stratified by MAIT cell subset **(G)**.
