## Supplemental Table2 for "CD4^+^ Mucosal-associated Invariant T (MAIT) cells express highly diverse T cell receptors"

**Supplemental Table 2:** Differential Gene expression between CD8 and CD4 MAIT cells

| | p_val | avg_log2FC | pct.1 | pct.2 | p_val_adj | genes |
|---|---|---|---|---|---|---|
| TRAV1-2 | 8.90E-57 | 4.588588 | 0.868 | 0.167 | 3.26E-52 | TRAV1-2 |
| SLC4A10 | 3.04E-47 | 3.738179 | 0.813 | 0.179 | 1.11E-42 | SLC4A10 |
| CTSW | 2.99E-40 | 1.846581 | 0.959 | 0.643 | 1.09E-35 | CTSW |
| KLRB1 | 6.31E-40 | 1.847105 | 0.964 | 0.53 | 2.31E-35 | KLRB1 |
| NKG7 | 1.41E-35 | 1.551821 | 0.976 | 0.732 | 5.16E-31 | NKG7 |
| NCR3 | 2.24E-35 | 2.19454 | 0.856 | 0.435 | 8.20E-31 | NCR3 |
| PRF1 | 3.19E-35 | 1.788724 | 0.935 | 0.583 | 1.17E-30 | PRF1 |
| MAL | 1.20E-34 | -2.40856 | 0.23 | 0.726 | 4.41E-30 | MAL |
| COTL1 | 2.16E-30 | -1.48587 | 0.746 | 0.952 | 7.91E-26 | COTL1 |
| CD4 | 9.53E-30 | -2.4261 | 0.113 | 0.542 | 3.49E-25 | CD4 |
| ZBTB16 | 2.71E-28 | 2.849718 | 0.669 | 0.173 | 9.92E-24 | ZBTB16 |
| TRBV6-4 | 6.34E-27 | 4.815175 | 0.535 | 0.054 | 2.32E-22 | TRBV6-4 |
| LST1 | 6.73E-27 | 2.119348 | 0.803 | 0.429 | 2.46E-22 | LST1 |
| KLRG1 | 9.53E-27 | 2.803763 | 0.655 | 0.19 | 3.49E-22 | KLRG1 |
| AL136456.1 | 5.18E-25 | 2.010919 | 0.803 | 0.423 | 1.90E-20 | AL136456.1 |
| GZMA | 1.07E-23 | 1.019275 | 0.983 | 0.792 | 3.90E-19 | GZMA |
| HOPX | 2.07E-23 | 1.131374 | 0.945 | 0.69 | 7.58E-19 | HOPX |
| CST7 | 3.59E-23 | 1.097199 | 0.969 | 0.798 | 1.31E-18 | CST7 |
| CEBPD | 1.43E-22 | 2.603052 | 0.633 | 0.196 | 5.22E-18 | CEBPD |
| IKZF2 | 7.56E-22 | 2.379737 | 0.619 | 0.214 | 2.77E-17 | IKZF2 |
| CCL4 | 6.51E-21 | 2.162744 | 0.794 | 0.411 | 2.38E-16 | CCL4 |
| CCL3 | 1.09E-20 | 2.698654 | 0.724 | 0.363 | 3.97E-16 | CCL3 |
| CD8A | 4.18E-20 | 1.235775 | 0.787 | 0.333 | 1.53E-15 | CD8A |
| LAG3 | 5.89E-20 | 1.577714 | 0.837 | 0.601 | 2.15E-15 | LAG3 |
| PLEK | 7.21E-20 | 1.942859 | 0.691 | 0.31 | 2.64E-15 | PLEK |
| GNLY | 1.21E-19 | 1.666099 | 0.923 | 0.714 | 4.41E-15 | GNLY |
| ADAM12 | 1.26E-19 | 2.736983 | 0.53 | 0.131 | 4.61E-15 | ADAM12 |
| TNFRSF4 | 7.54E-18 | -1.83563 | 0.463 | 0.738 | 2.76E-13 | TNFRSF4 |
| ARID5B | 1.04E-17 | -1.52793 | 0.319 | 0.655 | 3.81E-13 | ARID5B |
| HPGD | 2.11E-17 | 1.131856 | 0.693 | 0.327 | 7.74E-13 | HPGD |
| GZMK | 3.62E-17 | 1.465812 | 0.787 | 0.423 | 1.32E-12 | GZMK |
| ZFP36L1 | 1.01E-16 | 1.196243 | 0.863 | 0.673 | 3.71E-12 | ZFP36L1 |
| NME2 | 1.83E-16 | -0.8022 | 0.871 | 0.94 | 6.71E-12 | NME2 |
| RPS2 | 2.33E-16 | -0.53702 | 1 | 1 | 8.52E-12 | RPS2 |
| LYAR | 2.98E-16 | 1.587704 | 0.705 | 0.381 | 1.09E-11 | LYAR |
| IL4I1 | 7.24E-16 | 1.613083 | 0.607 | 0.25 | 2.65E-11 | IL4I1 |
| GZMB | 9.02E-16 | 1.038248 | 0.921 | 0.72 | 3.30E-11 | GZMB |
| CD81 | 1.70E-15 | 0.776296 | 0.959 | 0.875 | 6.24E-11 | CD81 |
| RPS8 | 4.05E-15 | -0.44475 | 1 | 1 | 1.48E-10 | RPS8 |
| LDHB | 1.02E-14 | -0.63695 | 0.928 | 0.964 | 3.75E-10 | LDHB |
| LINC01871 | 1.30E-14 | 0.861828 | 0.923 | 0.744 | 4.77E-10 | LINC01871 |
| RPL8 | 3.83E-14 | -0.35869 | 1 | 1 | 1.40E-09 | RPL8 |

| | | | | | | |
|---|---|---|---|---|---|---|
| TMIGD2 | 4.38E-14 | 1.624688 | 0.559 | 0.214 | 1.60E-09 | TMIGD2 |
| CCR1 | 6.00E-14 | 1.85107 | 0.523 | 0.19 | 2.20E-09 | CCR1 |
| CCL5 | 8.08E-14 | 0.40291 | 0.971 | 0.732 | 2.96E-09 | CCL5 |
| CTSH | 1.14E-13 | 1.194026 | 0.729 | 0.423 | 4.16E-09 | CTSH |
| RPLP2 | 1.40E-13 | -0.36188 | 1 | 1 | 5.13E-09 | RPLP2 |
| GBP5 | 2.34E-13 | 0.753383 | 0.894 | 0.75 | 8.55E-09 | GBP5 |
| ID2 | 2.67E-13 | 0.839094 | 0.902 | 0.744 | 9.76E-09 | ID2 |
| STAT1 | 3.42E-13 | -1.05245 | 0.743 | 0.869 | 1.25E-08 | STAT1 |
| RORA | 3.62E-13 | 0.840625 | 0.935 | 0.768 | 1.33E-08 | RORA |
| SEC11C | 4.13E-13 | -1.1722 | 0.659 | 0.804 | 1.51E-08 | SEC11C |
| CORO1B | 5.28E-13 | -1.24422 | 0.511 | 0.738 | 1.93E-08 | CORO1B |
| RPS18 | 8.37E-13 | -0.45446 | 1 | 1 | 3.06E-08 | RPS18 |
| LYST | 1.83E-12 | 1.121799 | 0.794 | 0.548 | 6.68E-08 | LYST |
| LINC00299 | 2.00E-12 | 1.329578 | 0.624 | 0.333 | 7.30E-08 | LINC00299 |
| TMSB10 | 3.20E-12 | -0.54762 | 1 | 1 | 1.17E-07 | TMSB10 |
| TNF | 4.62E-12 | 1.384895 | 0.7 | 0.429 | 1.69E-07 | TNF |
| GYG1 | 9.39E-12 | 0.922433 | 0.837 | 0.667 | 3.44E-07 | GYG1 |
| RPL12 | 1.33E-11 | -0.50395 | 1 | 1 | 4.88E-07 | RPL12 |
| PLCB1 | 1.81E-11 | 1.375482 | 0.602 | 0.292 | 6.63E-07 | PLCB1 |
| CXCR6 | 1.83E-11 | 0.768224 | 0.868 | 0.577 | 6.68E-07 | CXCR6 |
| SIRPG | 2.51E-11 | 1.483811 | 0.561 | 0.268 | 9.18E-07 | SIRPG |
| PRDX1 | 3.69E-11 | -0.86747 | 0.878 | 0.946 | 1.35E-06 | PRDX1 |
| EEF2 | 5.04E-11 | -0.51783 | 0.935 | 0.988 | 1.85E-06 | EEF2 |
| BTF3 | 8.15E-11 | -0.38562 | 0.988 | 1 | 2.98E-06 | BTF3 |
| CD69 | 9.08E-11 | 0.747316 | 0.916 | 0.786 | 3.32E-06 | CD69 |
| LY6E | 9.42E-11 | -0.74948 | 0.863 | 0.935 | 3.45E-06 | LY6E |
| TMEM173 | 1.18E-10 | -1.32622 | 0.412 | 0.655 | 4.32E-06 | TMEM173 |
| RPS16 | 1.68E-10 | -0.36996 | 1 | 1 | 6.14E-06 | RPS16 |
| RPS6 | 2.06E-10 | -0.40762 | 1 | 1 | 7.54E-06 | RPS6 |
| PARP8 | 2.20E-10 | 0.966744 | 0.791 | 0.571 | 8.04E-06 | PARP8 |
| TXK | 2.60E-10 | 1.159449 | 0.633 | 0.375 | 9.50E-06 | TXK |
| KIAA0319L | 3.75E-10 | 1.113741 | 0.652 | 0.429 | 1.37E-05 | KIAA0319L |
| GSTP1 | 5.00E-10 | -0.68559 | 0.787 | 0.905 | 1.83E-05 | GSTP1 |
| RBM3 | 7.71E-10 | -0.61388 | 0.818 | 0.887 | 2.82E-05 | RBM3 |
| CMTM7 | 9.14E-10 | -1.33877 | 0.264 | 0.506 | 3.35E-05 | CMTM7 |
| RPL18A | 9.27E-10 | -0.35726 | 1 | 1 | 3.39E-05 | RPL18A |
| LINC01943 | 1.03E-09 | -1.20997 | 0.446 | 0.673 | 3.79E-05 | LINC01943 |
| PTMS | 1.12E-09 | 0.937632 | 0.767 | 0.595 | 4.11E-05 | PTMS |
| RPL19 | 1.22E-09 | -0.35328 | 1 | 1 | 4.46E-05 | RPL19 |
| LTB | 1.26E-09 | 0.561497 | 0.998 | 1 | 4.60E-05 | LTB |
| TSPO | 1.32E-09 | -0.73555 | 0.647 | 0.821 | 4.85E-05 | TSPO |
| SYTL2 | 1.62E-09 | 0.778156 | 0.758 | 0.524 | 5.95E-05 | SYTL2 |
| SUPT3H | 1.89E-09 | 1.002707 | 0.621 | 0.363 | 6.92E-05 | SUPT3H |
| GBP1 | 2.27E-09 | -0.99969 | 0.338 | 0.577 | 8.32E-05 | GBP1 |
| SUB1 | 3.17E-09 | -0.51403 | 0.952 | 0.988 | 0.000116 | SUB1 |
| APOBEC3G | 3.35E-09 | 0.769494 | 0.803 | 0.667 | 0.000122 | APOBEC3G |

| | | | | | | |
|---|---|---|---|---|---|---|
| PHACTR2 | 3.36E-09 | 1.051561 | 0.693 | 0.518 | 0.000123 | PHACTR2 |
| RPSA | 3.84E-09 | -0.38285 | 1 | 1 | 0.000141 | RPSA |
| ALOX5AP | 4.14E-09 | 0.504623 | 0.89 | 0.702 | 0.000151 | ALOX5AP |
| CCNG2 | 5.08E-09 | 1.273223 | 0.513 | 0.25 | 0.000186 | CCNG2 |
| RPL4 | 5.81E-09 | -0.46802 | 0.94 | 0.982 | 0.000213 | RPL4 |
| PITPNC1 | 5.96E-09 | 0.931527 | 0.734 | 0.5 | 0.000218 | PITPNC1 |
| RPL10A | 7.77E-09 | -0.35924 | 0.99 | 0.994 | 0.000284 | RPL10A |
| CLEC2B | 8.02E-09 | 0.581997 | 0.868 | 0.673 | 0.000294 | CLEC2B |
| HMGB2 | 8.84E-09 | 1.317762 | 0.779 | 0.708 | 0.000324 | HMGB2 |
| ARL6IP5 | 1.09E-08 | -0.48978 | 0.978 | 0.988 | 0.0004 | ARL6IP5 |
| ETFB | 1.24E-08 | -0.90845 | 0.518 | 0.685 | 0.000455 | ETFB |
| MALAT1 | 1.26E-08 | 0.441939 | 1 | 1 | 0.000462 | MALAT1 |
| RPL22 | 1.28E-08 | -0.37728 | 0.986 | 1 | 0.000467 | RPL22 |
| TIMP1 | 1.30E-08 | -1.1432 | 0.338 | 0.565 | 0.000475 | TIMP1 |
| RUNX2 | 1.61E-08 | 1.002096 | 0.595 | 0.327 | 0.00059 | RUNX2 |
| NPM1 | 2.05E-08 | -0.46062 | 0.964 | 0.994 | 0.000752 | NPM1 |
| MKI67 | 2.18E-08 | 1.746633 | 0.523 | 0.315 | 0.000799 | MKI67 |
| RPS11 | 2.20E-08 | -0.39957 | 0.976 | 1 | 0.000805 | RPS11 |
| RGS1 | 2.32E-08 | -1.23533 | 0.353 | 0.571 | 0.00085 | RGS1 |
| RPL5 | 2.51E-08 | -0.40822 | 0.99 | 0.988 | 0.000918 | RPL5 |
| PIP4K2A | 2.62E-08 | 0.840976 | 0.739 | 0.542 | 0.000957 | PIP4K2A |
| SMAP2 | 2.87E-08 | -0.7363 | 0.703 | 0.833 | 0.00105 | SMAP2 |
| ARL3 | 3.74E-08 | 1.009834 | 0.561 | 0.315 | 0.001371 | ARL3 |
| COX6B1 | 4.18E-08 | -0.36224 | 0.957 | 0.994 | 0.001532 | COX6B1 |
| LAT | 6.11E-08 | -0.70832 | 0.775 | 0.845 | 0.002235 | LAT |
| DDIT4 | 7.17E-08 | -1.16669 | 0.616 | 0.768 | 0.002625 | DDIT4 |
| DNAJC1 | 1.01E-07 | 0.753641 | 0.77 | 0.631 | 0.00368 | DNAJC1 |
| CELF2 | 1.05E-07 | 0.623962 | 0.918 | 0.833 | 0.003844 | CELF2 |
| PTPN22 | 1.19E-07 | 0.677932 | 0.753 | 0.589 | 0.004362 | PTPN22 |
| SUSD3 | 1.25E-07 | -1.0229 | 0.305 | 0.512 | 0.004581 | SUSD3 |
| LCP1 | 1.34E-07 | 0.439917 | 0.964 | 0.905 | 0.004916 | LCP1 |
| ISG20 | 1.47E-07 | -1.12747 | 0.528 | 0.69 | 0.005384 | ISG20 |
| PSMA2 | 1.66E-07 | -0.45478 | 0.813 | 0.917 | 0.006063 | PSMA2 |
| PHLDA1 | 1.68E-07 | 0.931302 | 0.621 | 0.405 | 0.006162 | PHLDA1 |
| HCST | 2.11E-07 | 0.422512 | 0.988 | 0.964 | 0.007718 | HCST |
| GAB3 | 2.17E-07 | 1.021031 | 0.501 | 0.274 | 0.007931 | GAB3 |
| EEF1G | 3.13E-07 | -0.36926 | 0.969 | 0.988 | 0.011457 | EEF1G |
| TRAPPC1 | 3.45E-07 | -0.40376 | 0.923 | 0.923 | 0.012624 | TRAPPC1 |
| HSPD1 | 3.49E-07 | -0.74961 | 0.65 | 0.78 | 0.012761 | HSPD1 |
| FKBP1A | 3.70E-07 | -0.57144 | 0.779 | 0.863 | 0.013552 | FKBP1A |
| MYO1F | 3.92E-07 | 0.580405 | 0.791 | 0.607 | 0.014365 | MYO1F |
| CBLB | 4.06E-07 | 0.662562 | 0.791 | 0.643 | 0.01487 | CBLB |
| ITGB2 | 4.09E-07 | 0.559678 | 0.892 | 0.792 | 0.014961 | ITGB2 |
| DDX39A | 5.09E-07 | 0.886913 | 0.655 | 0.476 | 0.01863 | DDX39A |
| RPL9 | 5.90E-07 | -0.40578 | 0.998 | 1 | 0.021584 | RPL9 |
| SLC25A3 | 5.96E-07 | -0.4494 | 0.866 | 0.917 | 0.021802 | SLC25A3 |

| | | | | | | |
|---|---|---|---|---|---|---|
| IL2RA | 6.01E-07 | -0.90817 | 0.604 | 0.762 | 0.021992 | IL2RA |
| TNFRSF1B | 6.03E-07 | -0.78365 | 0.76 | 0.821 | 0.022086 | TNFRSF1B |
| TRG-AS1 | 6.21E-07 | 0.828521 | 0.65 | 0.464 | 0.022745 | TRG-AS1 |
| TUBA1A | 6.32E-07 | 0.825737 | 0.679 | 0.524 | 0.023126 | TUBA1A |
| GSTK1 | 6.97E-07 | -0.46684 | 0.916 | 0.935 | 0.025505 | GSTK1 |
| ENTPD1 | 7.23E-07 | 0.791855 | 0.7 | 0.565 | 0.026467 | ENTPD1 |
| FYB1 | 7.56E-07 | -0.66052 | 0.751 | 0.875 | 0.027685 | FYB1 |
| ATP8B4 | 7.78E-07 | 1.059799 | 0.58 | 0.375 | 0.028477 | ATP8B4 |
| ANK3 | 9.25E-07 | -0.97908 | 0.336 | 0.53 | 0.033865 | ANK3 |
| EFHD2 | 1.22E-06 | 0.518594 | 0.868 | 0.792 | 0.044731 | EFHD2 |
| MAPRE2 | 1.23E-06 | 0.856832 | 0.566 | 0.357 | 0.045055 | MAPRE2 |
| CKLF | 1.32E-06 | 0.374091 | 0.974 | 0.929 | 0.04822 | CKLF |
| LINC01138 | 1.50E-06 | 0.73088 | 0.679 | 0.482 | 0.054934 | LINC01138 |
| CCDC107 | 1.58E-06 | 0.599097 | 0.779 | 0.619 | 0.058004 | CCDC107 |
| S100A11 | 1.64E-06 | -0.36144 | 0.988 | 0.976 | 0.060085 | S100A11 |
| SOS1 | 1.74E-06 | -0.60864 | 0.76 | 0.851 | 0.063714 | SOS1 |
| NDUFS5 | 1.75E-06 | -0.43506 | 0.868 | 0.911 | 0.064086 | NDUFS5 |
| KLRK1 | 2.04E-06 | 0.789879 | 0.635 | 0.423 | 0.074582 | KLRK1 |
| VDAC1 | 2.07E-06 | -0.5121 | 0.741 | 0.815 | 0.075672 | VDAC1 |
| EEF1B2 | 2.30E-06 | -0.46084 | 0.957 | 0.994 | 0.084285 | EEF1B2 |
| TUBA1B | 2.34E-06 | 1.526324 | 0.813 | 0.756 | 0.085714 | TUBA1B |
| RNF157 | 2.39E-06 | 0.875367 | 0.59 | 0.399 | 0.087489 | RNF157 |
| PABPC1 | 2.60E-06 | -0.46388 | 0.918 | 0.97 | 0.095321 | PABPC1 |
| ACTR3 | 2.67E-06 | -0.45367 | 0.873 | 0.946 | 0.097778 | ACTR3 |
| NFKBIA | 2.90E-06 | 0.445794 | 0.918 | 0.845 | 0.106118 | NFKBIA |
| PSMB4 | 3.26E-06 | -0.61798 | 0.652 | 0.762 | 0.119439 | PSMB4 |
| RPS6KA3 | 3.35E-06 | 0.662391 | 0.767 | 0.601 | 0.122438 | RPS6KA3 |
| DPP4 | 3.58E-06 | 0.656536 | 0.657 | 0.452 | 0.130974 | DPP4 |
| COX5B | 3.85E-06 | -0.36956 | 0.926 | 0.946 | 0.140946 | COX5B |
| LMAN1 | 4.04E-06 | -0.61345 | 0.638 | 0.786 | 0.147696 | LMAN1 |
| BCL7C | 4.40E-06 | 0.676105 | 0.691 | 0.518 | 0.161178 | BCL7C |
| JAK1 | 4.42E-06 | 0.385853 | 0.942 | 0.899 | 0.161923 | JAK1 |
| PRR13 | 4.45E-06 | -0.3605 | 0.885 | 0.935 | 0.162809 | PRR13 |
| UBE2S | 4.82E-06 | 1.345029 | 0.609 | 0.5 | 0.176319 | UBE2S |
| TUBB | 5.34E-06 | 1.306603 | 0.868 | 0.839 | 0.195413 | TUBB |
| TGFBR2 | 5.68E-06 | 0.756435 | 0.643 | 0.452 | 0.20775 | TGFBR2 |
| CISH | 5.71E-06 | -0.88656 | 0.441 | 0.583 | 0.20907 | CISH |
| CTSD | 5.80E-06 | 0.519443 | 0.861 | 0.726 | 0.212137 | CTSD |
| FAM3C | 5.89E-06 | 0.86022 | 0.54 | 0.327 | 0.215505 | FAM3C |
| TAGLN2 | 6.09E-06 | -0.44035 | 0.911 | 0.94 | 0.222943 | TAGLN2 |
| MKNK2 | 6.11E-06 | -0.8801 | 0.384 | 0.565 | 0.223689 | MKNK2 |
| CCND2 | 6.20E-06 | -0.88518 | 0.71 | 0.804 | 0.227008 | CCND2 |
| CARS | 6.52E-06 | -0.95703 | 0.355 | 0.518 | 0.23849 | CARS |
| RPS17 | 6.52E-06 | -0.52034 | 0.866 | 0.887 | 0.23877 | RPS17 |
| CDC42EP3 | 6.92E-06 | 0.761634 | 0.58 | 0.375 | 0.253245 | CDC42EP3 |
| SAMD3 | 7.15E-06 | 0.796737 | 0.588 | 0.405 | 0.261614 | SAMD3 |

| | | | | | | |
|---|---|---|---|---|---|---|
| MDH2 | 7.40E-06 | -0.51403 | 0.7 | 0.798 | 0.270876 | MDH2 |
| C1QBP | 7.61E-06 | -0.76062 | 0.511 | 0.637 | 0.278523 | C1QBP |
| H2AFV | 8.09E-06 | 0.551382 | 0.897 | 0.875 | 0.296121 | H2AFV |
| TMPO | 8.44E-06 | 1.015891 | 0.607 | 0.488 | 0.309035 | TMPO |
| HP1BP3 | 8.79E-06 | 0.535376 | 0.82 | 0.685 | 0.321577 | HP1BP3 |
| TRAF3IP3 | 8.80E-06 | -0.42343 | 0.82 | 0.887 | 0.322065 | TRAF3IP3 |
| ARL6IP1 | 8.87E-06 | 0.742887 | 0.918 | 0.905 | 0.324507 | ARL6IP1 |
| PPA1 | 9.83E-06 | -0.81697 | 0.54 | 0.649 | 0.359642 | PPA1 |
| PYHIN1 | 1.03E-05 | 0.55153 | 0.719 | 0.536 | 0.375297 | PYHIN1 |
| NAP1L4 | 1.14E-05 | -0.52469 | 0.794 | 0.899 | 0.417065 | NAP1L4 |
| SSBP1 | 1.35E-05 | -0.47799 | 0.71 | 0.792 | 0.494775 | SSBP1 |
| H1FX | 1.40E-05 | 0.537649 | 0.902 | 0.857 | 0.513615 | H1FX |
| CHD9 | 1.44E-05 | 0.824993 | 0.535 | 0.351 | 0.525353 | CHD9 |
| FAF1 | 1.47E-05 | 0.756253 | 0.633 | 0.47 | 0.538365 | FAF1 |
| TAGAP | 1.47E-05 | 0.845765 | 0.561 | 0.381 | 0.538858 | TAGAP |
| NUCB2 | 1.52E-05 | 0.830041 | 0.547 | 0.369 | 0.556214 | NUCB2 |
| SNU13 | 1.64E-05 | -0.48674 | 0.794 | 0.851 | 0.602026 | SNU13 |
| FAM174C | 1.66E-05 | -0.67162 | 0.429 | 0.601 | 0.60847 | FAM174C |
| SIPA1L1 | 1.80E-05 | 0.91222 | 0.585 | 0.411 | 0.658157 | SIPA1L1 |
| IER5 | 1.96E-05 | 0.599805 | 0.614 | 0.423 | 0.717293 | IER5 |
| NCALD | 1.97E-05 | 0.515458 | 0.676 | 0.494 | 0.719834 | NCALD |
| COX14 | 1.99E-05 | -0.58258 | 0.604 | 0.756 | 0.728862 | COX14 |
| FKBP11 | 2.04E-05 | 0.468904 | 0.861 | 0.756 | 0.746837 | FKBP11 |
| SMC4 | 2.04E-05 | 1.163487 | 0.549 | 0.446 | 0.747089 | SMC4 |
| RNASET2 | 2.11E-05 | -0.79365 | 0.434 | 0.583 | 0.770535 | RNASET2 |
| STMN1 | 2.23E-05 | 1.104935 | 0.693 | 0.601 | 0.816626 | STMN1 |
| EVA1B | 2.31E-05 | 0.795181 | 0.513 | 0.345 | 0.847104 | EVA1B |
| TALDO1 | 2.34E-05 | -0.55171 | 0.763 | 0.821 | 0.855965 | TALDO1 |
| MRPS6 | 2.36E-05 | 0.446627 | 0.791 | 0.696 | 0.863321 | MRPS6 |
| TUBA4A | 2.41E-05 | 0.726114 | 0.727 | 0.637 | 0.880547 | TUBA4A |
| OCIAD2 | 2.41E-05 | -0.53078 | 0.715 | 0.798 | 0.882847 | OCIAD2 |
| UBE2F | 2.52E-05 | -0.92164 | 0.405 | 0.554 | 0.920923 | UBE2F |
| SLFN12L | 2.53E-05 | 0.625987 | 0.578 | 0.399 | 0.927476 | SLFN12L |
| DLEU2 | 2.62E-05 | 0.702018 | 0.612 | 0.452 | 0.960417 | DLEU2 |
| BUB3 | 2.71E-05 | 0.602424 | 0.775 | 0.685 | 0.993532 | BUB3 |
