## Supplemental Table3 for "CD4^+^ Mucosal-associated Invariant T (MAIT) cells express highly diverse T cell receptors"

**Supplemental Table 3:** CDR3α/β sequence similarity with published CDR3 sequences of T cells.

| **TRAV1-2 negative CDR3alpha sequence similarity** | | | | | | | | | | | | | | | | | |
|---|---|---|---|---|---|---|---|---|---|---|---|---|---|---|---|---|---|
| Query | Dist | CDR3.alpha.aa | Pathology | Pathology.Mesh.ID | Single.cell | Antigen.protein | Epitope.peptide | Epitope.ID | Tissue | T.Cell.Type | T.cell.characteristics | TRAV | TRAJ | TRBV | TRBD | TRBJ | PubMed.ID |
| CAVARSGGYQKVTF | 2 | CAASGGYQKVTF | Human immunodeficiency virus (HIV) | D006782 | No | Gag p24 | KAFSPEVIPMF | 29804 | PBMC | CD8 | NA | TRAV5 | NA | TRBV19 | NA | TRBJ 1-2 | 17121793 |
| CAEPGRRALTF | 2 | CAVSEPGRRALTF | M.Tuberculosis | D009169 | Yes | NA | NA | NA | PBMC | CD4 | NA | TRAV8-4 | TRAJ5 | TRBV29-1 | NA | TRBJ 2-5 | 32341563 |
| CAMSAGTGNQFYF | 2 | CAASGGTGNQFYF | Influenza | D009980 | Yes | Matrix protein (M1) | GILGFVFTL | 20354 | Bronchoalveolar | CD8 | NA | TRAV29/DV5:01 | TRAJ4-01 | TRBV1-01 | NA | TRBJ 2-7:01 | 28636592 |
| CLVRFGGFKTIF | 2 | CAVRGGGFKTIF | Parkinson disease | D010301 | Yes | NA | NA | NA | CSF | CD8 | NA | NA | TRAJ9 | TRBV2 | NA | TRBJ 2-3 | 31915375 |
| CAENNAGKSTF | 2 | CAVNAGKSTF | Yellow fever virus | D015005 | Yes | YFV-17D | LLWNGPMAV | 121572 | NA | CD8 | NA | TRAV12-2 | TRAJ27 | TRBV6-06 | TRBD1, TRBD2 | TRBJ 2-7 | 28103239 |
| CATDKAGTALIF | 1 | CATDAKAGTALIF | Human immunodeficiency virus (HIV) | D006678 | No | RF10 Protein Nef | RYPLTFGWCF | 56620 | PBMC | CD8 | NA | TRAV17 | TRAJ15 | TRBV28 | TRBD2 | TRBJ 2-2 | 24899498 |
| CAVRDRDGGFKTIF | 2 | CAVRDDTGGFKTIF | Melanoma | D008545 | Yes | GP100-IMD | IMDQVPFSV | 27469 | NA | NA | NA | NA | NA | TRBV19 | NA | NA | 30418433 |
| CALDMMDSNYQLIW | 2 | CAADMDSNYQLIW | Influenza | D009980 | Yes | PB1 | LSLRNPILV | 39494 | Bronchoalveolar | CD8 | NA | TRAV5N-4:01 | TRAJ3-01 | TRBV1-01 | NA | TRBJ 1-1:01 | 28636592 |
| CAVSGGGFKTIF | 2 | CAVNTGGFKTIF | Melanoma | D008545 | No | Melan-A/MART-1 | EAAGIGILTV | 10987 | Tumor Tissue | NA | NA | TRAV2 | TRAJ9 | NA | NA | NA | 7777568 |
| CAFSTAFGNEKLTF | 2 | CAFSTAAGNKLTF | Neoantigen | C050269 | Yes | BAIAP3-ILN | ILNVDVFTL | 606794 | NA | NA | NA | TRAV5 | NA | TRBV30 | NA | NA | 30418433 |
| CAVTNQAGTALIF | 2 | CAVVDQAGTALIF | Cytomegalovirus (CMV) | D003586 | Yes | pp65 | NLVPMVATV | 44920 | PBMC | CD8 | NA | TRAV2-01 | TRAJ1-01 | TRBV4-01 | NA | TRBJ 2-1:01 | 28636592 |

| | | | | | | | | | | | | | | | | | |
|---|---|---|---|---|---|---|---|---|---|---|---|---|---|---|---|---|---|
| CAVTLVW | 2 | CAVELVF | Neoantigen | C050269 | Yes | EXOC3L4-ILL | ILLDWAANV | NA | NA | NA | NA | TRAV14/DV4 | NA | TRBV7-9 | NA | NA | 30418433 |
| CARRSGGGADGLTF | 2 | CAVNSGGGADGLTF | Influenza | D009980 | Yes | Matrix protein (M1) | GILGFVFTL | 20354 | PBMC | CD8 | NA | TRAV21 | NA | TRBV6-1 | NA | NA | 28636589 |
| CAANSSASKIIF | 2 | CAEYSSASKIIF | Epstein Barr virus (EBV) | D020031 | No | BMLF-1 | GLCTLVAML | 20788 | PBMC | CD8 | NA | TRAV15 | TRAJ3 | TRBV14 | NA | TRBJ2-3 | 10925283 |
| CASRRPNTGNQFYF | 2 | CAYRSPNTGNQFYF | M.Tuberculosis | D009169 | Yes | NA | NA | NA | PBMC | CD4 | NA | mTRDV2-2 | TRAJ49 | TRBV11-3 | NA | TRBJ2-1 | 32341563 |
| CAFMKPNYGGATNKLIF | 2 | CAFLPPNYGGATNKLIF | M.Tuberculosis | D009169 | Yes | NA | NA | NA | PBMC | CD4 | NA | mTRDV2-2 | TRAJ32 | TRBV5-1 | NA | TRBJ1-2 | 32341563 |
| CAVVSYSGGGADGLTF | 2 | CAVPYSGGGADGLTF | Influenza | D009980 | Yes | Matrix protein (M1) | GILGFVFTL | 20354 | PBMC | CD8 | NA | TRAV12-2:01 | TRAJ4-01 | TRBV2-01 | NA | TRBJ2-5:01 | 28636592 |
| CAVSVAGGGADGLTF | 2 | CAVSRGGGADGLTF | Melanoma | D008545 | No | Melan-A/MART-1 | EAAGIGILTV | 10987 | Tumor Tissue | NA | NA | TRAV2 | TRAJ45 | NA | NA | TRBJ1-5 | 7777568 |
| CATASSGNTPLVF | 2 | CAETPSSGNTPLVF | Neoantigen | C050269 | Yes | OR6F1-VLN | VLNPFIYTL | NA | NA | NA | NA | TRAV12-2 | NA | TRBV4-1 | NA | NA | 30418433 |
| CAVNVTGGGNKLTF | 2 | CAVIFTGGGNKLTF | Human immunodeficiency virus (HIV) | D006678 | No | RF10 Protein Nef | RYPLTFGWCF | 56620 | PBMC | CD8 | NA | TRAV8-1 | TRAJ10 | TRBV7-09 | TRBD1 | TRBJ2-5 | 24899498 |
| CAVSDTNAGKSTF | 2 | CAVRDTNTGKSTF | Herpes simplex virus 2 (HSV2) | D018258 | No | HSV-2 VP22 | RPRGEVRFL | 55320 | PBMC | CD8 | NA | TRAV19 | NA | TRBV9 | NA | TRBJ2-1 | 20139278 |
| CALKRQGGKLIF | 2 | CALSNQGGKLIF | Influenza | D009980 | Yes | Matrix protein (M1) | GILGFVFTL | 20354 | PBMC | CD8 | NA | TRAV19 | NA | TRBV19 | NA | NA | 28636589 |
| CAYRSARVKTSGSRLTF | 2 | CAYRSARTSGSRLTF | M.Tuberculosis | D009169 | Yes | NA | NA | NA | PBMC | CD4 | NA | mTRDV2-2 | TRAJ58 | TRBV7-2 | NA | TRBJ1-4 | 32341563 |
| CAESRMAAGNKLTF | 2 | CAASMAAGNKLTF | HTLV-1 | D015368 | No | NA | NA | NA | PBMC | CD4 | NA | NA | TRAJ17 | NA | TRBD1 | TRBJ2-3 | 31069115 |
| CAVQLGENAGNMLTF | 2 | CAVQGANAGNMLTF | M.Tuberculosis | D009169 | Yes | NA | NA | NA | PBMC | CD4 | NA | TRAV2 | TRAJ39 | TRBV4-1 | NA | TRBJ1-4 | 32341563 |

| | | | | | | | | | | | | | | | | | |
|---|---|---|---|---|---|---|---|---|---|---|---|---|---|---|---|---|---|
| CVVRALSGAGSYQLTF | 2 | CVVVAYSGAGSYQLTF | Cytomegalovirus (CMV) | D003586 | Yes | pp65 | NLVPMVATV | 44920 | PBMC | CD8 | NA | TRAV10 | NA | TRBV7-2 | NA | NA | 28636589 |
| CASSYNTDKLIF | 2 | CAVSGSYNTDKLIF | Cytomegalovirus (CMV) | D003586 | Yes | pp65 | NLVPMVATV | 44920 | PBMC | CD8 | NA | TRAV8-4:01 | TRAJ3-01 | TRBV2-01 | NA | TRBJ 2-7:01 | 28636592 |
| CVVRVTGGGNKLTF | 2 | CAVRRTGGGNKLTF | M.Tuberculosis | D009169 | Yes | NA | NA | NA | PBMC | CD4 | NA | TRAV3 | TRAJ10 | TRBV3-1 | NA | TRBJ 1-2 | 32341563 |
| CAASAPGGTSYGKLTF | 1 | CAASASGGTSYGKLTF | Influenza | D009980 | Yes | Matrix protein (M1) | GILGFVFTL | 20354 | PBMC | CD8 | NA | NA | NA | TRBV6-5:01 | TRBD | TRBJ 2-3:01 | 28300170 |
| CAVGVVGSQGNLIF | 2 | CAVGGSQGNLIF | Influenza | D009980 | Yes | Matrix protein (M1) | GILGFVFTL | 20354 | Bronchoalveolar | CD8 | NA | TRAV8-6:02 | TRAJ4-01 | TRBV1-01 | NA | TRBJ 2-7:01 | 28636592 |
| CATGDSKLTF | 2 | CAGGDSTLTF | Yellow fever virus | D015005 | Yes | YFV-17D | LLWNGPMAV | 121572 | NA | CD8 | NA | TRAV12-2 | TRAJ11 | TRBV3-01 | TRBD1, TRBD2 | TRBJ 1-5 | 28103239 |
| CAVNKAAGNKLTF | 2 | CVVFKAAGNKLTF | Epstein Barr virus (EBV) | D020031 | Yes | BMLF-1 | GLCTLVAML | 20788 | PBMC | CD8 | NA | TRAV10-01 | TRAJ1-01 | TRBV1-01 | NA | TRBJ 2-5:01 | 28636592 |
| CAMREVNSGGYQKVTF | 2 | CAMRDLNSGGYQKVTF | Neoantigen | C050269 | Yes | PGM5-AVG-H5Y | AVGSYVYSV | NA | NA | NA | NA | TRAV12-2 | NA | TRBV11-2 | NA | NA | 30418433 |
| CAVEENDYKLSF | 2 | CAVSSNDYKLSF | Yellow fever virus | D015005 | Yes | YFV-17D | LLWNGPMAV | 121572 | NA | CD8 | NA | TRAV12-2 | TRAJ20 | TRBV6-01 | TRBD2 | TRBJ 1-5 | 28103239 |
| CAVSLNRDDKIIF | 2 | CAVRNRDDKIIF | Neoantigen | C050269 | Yes | WDR46 | FLTYLDVSV | NA | NA | NA | NA | NA | NA | TRBV9 | NA | NA | 30418433 |
| CAVPHNFGNEKLTF | 2 | CAVRANFGNEKLTF | Influenza | D009980 | Yes | Matrix protein (M1) | GILGFVFTL | 20354 | PBMC | CD8 | NA | TRAV41:01 | TRAJ4-01 | TRBV6-01 | NA | TRBJ 2-2:01 | 28636592 |
| CAMSERGFGNVLHC | 2 | CAVSEGFGNVLHC | M. tuberculosis | D009169 | Yes | NA | NA | NA | PBMC | CD8 | NA | NA | TRAJ35 | TRBV20-1 | NA | TRBJ 2-7 | 30992377 |
| CATHNTDKLIF | 2 | CAALNTDKLIF | Diabetes Type 1 | D003922 | No | GAD65 | NFIRMVISNPAAT | 101069 | PBMC | CD4 | NA | NA | NA | NA | NA | TRBJ 1-1 | 25681349 |
| CAVGSAGGTSYGKLTF | 2 | CAMSSAGGTSYGKLTF | Epstein Barr virus (EBV) | D020031 | Yes | BMLF-1 | GLCTLVAML | 20788 | PBMC | CD8 | NA | TRAV12-3 | NA | TRBV14 | NA | NA | 28636589 |
| CAVNGLGFGNVLHC | 2 | CAVEDLGFGNVLHC | Neoantigen | C050269 | Yes | SEC24A | FLYNPLTRV | NA | NA | NA | NA | TRAV12-3 | NA | TRBV28 | NA | NA | 30418433 |

| | | | | | | | | | | | | | | | | | |
|---|---|---|---|---|---|---|---|---|---|---|---|---|---|---|---|---|---|
| CAVNVVGTSYGKLTF | 2 | CAVNAGGTSYGKLTF | Diabetes Type 1 | D003922 | Yes | NA | NA | NA | Pancreatic islets | CD4 | NA | TRAV12-2 | TRAJ52 | TRBV6-5 | NA | TRBJ 1-1 | 27920090 |
| CAVGNQGGSEKLVF | 2 | CAGTQGGSEKLVF | Tumor | D009370 | No | NY-ESO-1 | NA | NA | PBL | CD4 | NA | TRAV38-1 | TRAJ57 | TRBV2 | NA | TRBJ 2-3 | 30530988 |
| CAVTPRANDYKLSF | 2 | CAVPRNDYKLSF | Neoantigen | C050269 | Yes | OR14C36-FML-V6L | FMLYLLTLM | NA | NA | NA | NA | TRAV19 | NA | TRBV5-1 | NA | NA | 30418433 |
| CAVRGSGSARQLTF | 2 | CAVSGSARQLTF | Yellow fever virus | D015005 | Yes | YFV-17D | LLWNGPMAV | 121572 | NA | CD8 | NA | TRAV12-2 | TRAJ22 | TRBV9 | TRBD1, TRBD2 | TRBJ 2-7 | 28103239 |
| CATDASGYSTLTF | 2 | CATLSGYSTLTF | M.Tuberculosis | D009169 | Yes | NA | NA | NA | PBMC | CD4 | NA | TRAV17 | TRAJ11 | TRBV25-1 | NA | TRBJ 2-2 | 32341563 |
| CACYDMNRDDKIIF | 2 | CADDMNRDDKIIF | Neoantigen | C050269 | Yes | FLNA-HIA | HIAKSPFEV | NA | NA | NA | NA | NA | NA | TRBV6-2,6-3 | NA | NA | 30418433 |
| CAVKRGNEKLTF | 2 | CAVTGNEKLTF | Yellow fever virus | D015005 | Yes | YFV-17D | LLWNGPMAV | 121572 | NA | CD8 | NA | TRAV12-2 | TRAJ48 | TRBV6-05 | TRBD2 | TRBJ 2-7 | 28103239 |
| CALTKASGGSNYKLTF | 2 | CALTQNSGGSNYKLTF | M.Tuberculosis | D009169 | Yes | NA | NA | NA | PBMC | CD4 | NA | TRAV9-1 | TRAJ53 | TRBV20-1 | NA | TRBJ 1-2 | 32341563 |
| CAVCNNNDMRF | 2 | CAVTRNNDMRF | Yellow fever virus | D015005 | Yes | YFV-17D | LLWNGPMAV | 121572 | NA | CD8 | NA | TRAV12-2 | TRAJ43 | TRBV5-04 | TRBD2 | TRBJ 2-7 | 28103239 |
| CALCPTGANSKLTF | 2 | CATGPTGANSKLTF | M.Tuberculosis | D009169 | Yes | NA | NA | NA | PBMC | CD4 | NA | TRAV17 | TRAJ56 | TRBV20-1 | NA | TRBJ 1-4 | 32341563 |
| CALSGGSNYKLTF | 2 | CAVSGGINYKLTF | Tumor | D009369 | No | T72(Tn) | VITAFTEGLK | 693671 | NA | NA | NA | TRAV1 | TRAJ45 | TRBV1 | NA | TRBJ 2-1 | 10508250 |
| CLLATYSGAGSYQLTF | 2 | CAATYSGAGSYQLTF | M. tuberculosis | D009169 | Yes | NA | NA | NA | PBMC | CD8 | NA | NA | TRAJ28 | TRBV2 | NA | TRBJ 2-3 | 30992377 |
| CAVSEWGGSEKLVF | 2 | CAVSSGGGSEKLVF | M.Tuberculosis | D009169 | Yes | NA | NA | NA | PBMC | CD4 | NA | TRAV21 | TRAJ57 | TRBV3-1 | NA | TRBJ 1-6 | 32341563 |
| CLVGSLSGGYNKLIF | 2 | CLVGDSGGYNKLIF | M.Tuberculosis | D009169 | Yes | NA | NA | NA | PBMC | CD4 | NA | TRAV4 | TRAJ4 | TRBV20-1 | NA | TRBJ 2-2 | 32341563 |
| CAFTTNTGNQFYF | 2 | CATKNTGNQFYF | Cytomegalovirus (CMV) | D003586 | No | pp65 | NLVPMVATV | 44920 | PBMC | CD8 | NA | TRAV2-2 | TRAJ49 | TRBV14 | NA | TRBJ 2-3 | 16237109 |
| CAVSSNRDDKIIF | 2 | CAVRNRDDKIIF | Neoantigen | C050269 | Yes | WDR46 | FLTYLDVSV | NA | NA | NA | NA | NA | NA | TRBV9 | NA | NA | 30418433 |
| CAVLYSGNTGKLIF | 2 | CAVDGSGNTGKLIF | Diabetes Type 1 | D003922 | Yes | NA | NA | NA | Pancreatic islets | CD4 | NA | TRAV2 | TRAJ37 | TRBV4-1 | NA | TRBJ 2-7 | 27920090 |

| | | | | | | | | | | | | | | | | | |
|---|---|---|---|---|---|---|---|---|---|---|---|---|---|---|---|---|---|
| CATDEGAQKLVF | 2 | CATDHQGAQKLVF | Yellow fever virus | D015005 | Yes | YFV-17D | LLWNGPMAV | 121572 | NA | CD8 | NA | TRAV17 | TRAJ54 | TRBV9 | TRBD2 | TRBJ2-1 | 28103239 |
| CASFRIQGAQKLVF | 2 | CAFIQGAQKLVF | Influenza | D009980 | Yes | Matrix protein (M1) | GILGFVFTL | 20354 | PBMC | CD8 | NA | TRAV38-1 | NA | TRBV27 | NA | NA | 28636589 |
| CAALPYSGGGADGLTF | 2 | CAVPYSGGGADGLTF | Influenza | D009980 | Yes | Matrix protein (M1) | GILGFVFTL | 20354 | PBMC | CD8 | NA | TRAV12-2:01 | TRAJ4-01 | TRBV2-01 | NA | TRBJ2-5:01 | 28636592 |
| CAATNTGTASKLTF | 2 | CALNTGTASKLTF | Neoantigen | C050269 | Yes | NSDHL-A9V | KLVALGINAV | NA | NA | NA | NA | TRAV38-2/DV8 | NA | TRBV7-8 | NA | NA | 30418433 |
| CANGGGADGLTF | 2 | CAVNSGGGADGLTF | Influenza | D009980 | Yes | Matrix protein (M1) | GILGFVFTL | 20354 | PBMC | CD8 | NA | TRAV21 | NA | TRBV6-1 | NA | NA | 28636589 |
| CAVIPSGGYQKVTF | 2 | CAVSGGYQKVTF | Human immunodeficiency virus (HIV) | D006783 | No | Gag p24 | KAFSPEVIPMF | 29804 | PBMC | CD8 | NA | TRAV5 | NA | TRBV19 | NA | TRBJ1-2 | 17121793 |
| CAMREGVTGNQFYF | 2 | CAMREGLQTGNQFYF | M. tuberculosis | D009169 | Yes | NA | NA | NA | PBMC | CD8 | NA | NA | TRAJ49 | TRBV6-1 | NA | TRBJ2-7 | 30992377 |
| CAMSEAAGNKLTF | 2 | CAFSTAAGNKLTF | Neoantigen | C050269 | Yes | BAIAP3-ILN | ILNVDVFTL | 606794 | NA | NA | NA | TRAV5 | NA | TRBV30 | NA | NA | 30418433 |
| CALVYSSASKIIF | 2 | CAEYSSASKIIF | Epstein Barr virus (EBV) | D020031 | No | BMLF-1 | GLCTLVAML | 20788 | PBMC | CD8 | NA | TRAV15 | TRAJ3 | TRBV14 | NA | TRBJ2-3 | 10925283 |
| CAFMRGAQKLVF | 1 | CAFGRGAQKLVF | Herpes simplex virus 2 (HSV2) | D018258 | No | HSV-2 VP22 | RPRGEVRFL | 55320 | PBMC | CD8 | NA | TRAV5 | NA | TRBV30 | NA | TRBJ1-4 | 20139278 |
| CAVNLQGAQKLVF | 2 | CAVIIQGAQKLVF | Tumor | D009370 | No | NY-ESO-1 | NA | NA | PBL | CD4 | NA | TRAV12-2 | TRAJ54 | TRBV2 | NA | TRBJ1-5 | 30530988 |
| CGAAFDSWGKLQF | 2 | CAATDSWGKLQF | Yellow fever virus | D015005 | Yes | YFV-17D | LLWNGPMAV | 121572 | NA | CD8 | NA | TRAV25 | TRAJ24 | TRBV9 | TRBD1, TRBD2 | TRBJ2-1 | 28103239 |
| CAVRYNNAGNMLTF | 2 | CAVREDNAGNMLTF | Cytomegalovirus (CMV) | D003586 | Yes | pp65 | NLVPMVATV | 44920 | PBMC | CD8 | NA | TRAV1-2:01 | TRAJ3-01 | TRBV1-01 | NA | TRBJ2-1:01 | 28636592 |
| CAVNAYNDMRF | 2 | CAVNMGNDMRF | Neoantigen | C050269 | Yes | MRM1-9 | LLFGMTPCL | NA | NA | NA | NA | NA | NA | TRBV4-2 | NA | NA | 30418433 |

| | | | | | | | | | | | | | | | | | |
|---|---|---|---|---|---|---|---|---|---|---|---|---|---|---|---|---|---|
| CAVSPPGSARQLTF | 2 | CAVSGSARQLTF | Yellow fever virus | D015005 | Yes | YFV-17D | LLWNGPMAV | 121572 | NA | CD8 | NA | TRAV12-2 | TRAJ22 | TRBV9 | TRBD1, TRBD2 | TRBJ2-7 | 28103239 |
| CVVRSYNFNKFYF | 2 | CVVIVYNFNKFYF | M.Tuberculosis | D009169 | Yes | NA | NA | NA | PBMC | CD4 | NA | TRAV8-4 | TRAJ21 | TRBV2 | NA | TRBJ2-7 | 32341563 |
| CATSGNTPLVF | 2 | CAVDSGNTPLVF | Yellow fever virus | D015005 | Yes | YFV-17D | LLWNGPMAV | 121572 | NA | CD8 | NA | TRAV12-2 | TRAJ29 | TRBV12-04 | TRBD1 | TRBJ2-7 | 28103239 |
| CVALSGTYKYIF | 2 | CAVSGTYKYIF | Yellow fever virus | D015005 | Yes | YFV-17D | LLWNGPMAV | 121572 | NA | CD8 | NA | TRAV12-2 | TRAJ40 | TRBV4-03 | TRBD2 | TRBJ2-1 | 28103239 |
| CAATDTGNQFYF | 2 | CAVADTGNQFYF | Cytomegalovirus (CMV) | D003586 | Yes | pp65 | NLVPMVATV | 44920 | PBMC | CD8 | NA | TRAV22:01 | TRAJ4-01 | TRBV1-01 | NA | TRBJ2-3:01 | 28636592 |
| CALASNTGNQFYF | 2 | CAENSNTGNQFYF | Epstein Barr virus (EBV) | D020031 | No | BMLF-1 | GLCTLVAML | 20788 | PBMC | CD8 | NA | TRAV15 | TRAJ49 | TRBV14 | NA | TRBJ2-3 | 10925283 |
| CVVNVGGGGNKLTF | 1 | CVVNLGGGGNKLTF | M.Tuberculosis | D009169 | Yes | NA | NA | NA | PBMC | CD4 | NA | TRAV12-1 | TRAJ10 | TRBV5-1 | NA | TRBJ1-4 | 32341563 |
| CVVTSNYGQNFVF | 2 | CAVTDNYGQNFVF | Neoantigen | C050269 | Yes | HAUS3-ILN-T7A | ILNAMIAKI | NA | NA | NA | NA | TRAV29/DV5 | NA | TRBV6-5 | NA | NA | 30418433 |
| CAVGDRNTGFQKLVF | 2 | CAVRDNTGFQKLVF | Diabetes Type 1 | D003922 | Yes | NA | NA | NA | Pancreatic islets | CD4 | NA | TRAV8-4 | TRAJ8 | TRBV6-1 | NA | TRBJ2-5 | 27920090 |
| CAPAGADKLIF | 2 | CAGADKLIF | Neoantigen | C050269 | Yes | GNL3L-R4C | NLNCCSVPV | NA | NA | NA | NA | TRAV3 | NA | TRBV6-5 | NA | NA | 30418433 |
| CAVLVSNFGNEKLTF | 2 | CAVSPVSNFGNEKLTF | Influenza | D009980 | Yes | Matrix protein (M1) | GILGFVFTL | 20354 | PBMC | CD8 | NA | TRAV8-6:02 | TRAJ4-01 | TRBV5-01 | NA | TRBJ2-1:01 | 28636592 |
| CASQRYGGSQGNLIF | 2 | CAVRYGGSQGNLIF | Breast Cancer | D001943 | No | NA | NA | NA | Breast | CD8 | TIL | TRAV41 | TRAJ42 | TRBV7-03 | NA | TRBJ2-4 | 27307436 |
| CVVSGSTYKYIF | 2 | CAVSGTYKYIF | Yellow fever virus | D015005 | Yes | YFV-17D | LLWNGPMAV | 121572 | NA | CD8 | NA | TRAV12-2 | TRAJ40 | TRBV4-03 | TRBD2 | TRBJ2-1 | 28103239 |
| CAVSSSGGSYIPTF | 2 | CAEISSGGSYIPTF | Influenza | D009980 | Yes | Matrix protein (M1) | GILGFVFTL | 20354 | PBMC | CD8 | NA | TRAV13-2:01 | TRAJ6-01 | TRBV9-01 | NA | TRBJ2-3:01 | 28636592 |
| CAARSYNTDKLIF | 2 | CAARANTDKLIF | Yellow fever virus | D015005 | Yes | YFV-17D | LLWNGPMAV | 121572 | NA | CD8 | NA | TRAV29/DV5 | TRAJ34 | TRBV15 | TRBD1 | TRBJ1-4 | 28103239 |
| CAYRSGLTGANSKLTF | 2 | CAYRSTTGANSKLTF | M.Tuberculosis | D009169 | Yes | NA | NA | NA | PBMC | CD4 | NA | mTRDV2-2 | TRAJ56 | TRBV20-1 | NA | TRBJ1-6 | 32341563 |

| | | | | | | | | | | | | | | | | | |
|---|---|---|---|---|---|---|---|---|---|---|---|---|---|---|---|---|---|
| CAGPSGGTYKYIF | 1 | CAGPSGTYKYIF | M.Tuberculosis | D009169 | Yes | NA | NA | NA | PBMC | CD4 | NA | TRAV35 | TRAJ40 | TRBV11-3 | NA | TRBJ2-5 | 32341563 |
| CAVIRTTGNQFYF | 2 | CAVRLTGNQFYF | M.Tuberculosis | D009169 | Yes | NA | NA | NA | PBMC | CD4 | NA | TRAV21 | TRAJ49 | TRBV29-1 | NA | TRBJ1-3 | 32341563 |
| CAVQRRGFQKLVF | 2 | CAVRRGAQKLVF | M.Tuberculosis | D009169 | Yes | NA | NA | NA | PBMC | CD4 | NA | TRAV21 | TRAJ54 | TRBV12-3 | NA | TRBJ1-6 | 32341563 |
| CAGRNSGGYQKVTF | 2 | CAGSGGYQKVTF | Diabetes Type 1 | D003922 | No | GAD65 | NA | NA | PBMC | CD4 | NA | NA | NA | NA | NA | TRBJ2-1 | 25681349 |
| CVVNGGGSNYKLTF | 2 | CAVGGGGSNYKLTF | M.Tuberculosis | D009169 | Yes | NA | NA | NA | PBMC | CD4 | NA | TRAV22 | TRAJ53 | TRBV6-1 | NA | TRBJ1-1 | 32341563 |
| CAMREANTNAGKSTF | 1 | CAMREAYNTNAGKSTF | Diabetes Type 1 | D003922 | Yes | NA | NA | NA | Pancreatic islets | CD4 | NA | TRDV1 | TRAJ19 | TRBV18 | NA | TRBJ1-3 | 27920090 |
| CLVGPAAGNKLTF | 1 | CLVGEAAGNKLTF | Celiac disease | D002507 | Yes | DQ-a-II | PQPELPYPQ | 49047 | Small intestine | CD4 | NA | TRAV12-2 | TRAJ17-1 | TRBV7-2*01 | TRBD2*01 | TRBJ2-7*01 | 24777060 |
| CAFHGSSNTGKLIF | 2 | CAGAHGSSNTGKLIF | Influenza | D009980 | Yes | Matrix protein (M1) | GILGFVFTL | 20354 | PBMC | CD8 | NA | TRAV2-01 | TRAJ3-01 | TRBV1-01 | NA | TRBJ2-7:01 | 28636592 |
| CAVRSEDSSYKLIF | 2 | CAVRATDSSYKLIF | Cytomegalovirus (CMV) | D003586 | Yes | pp65 | NLVPMVATV | 44920 | PBMC | CD8 | NA | TRAV1-2:01 | TRAJ1-01 | TRBV7-01 | NA | TRBJ2-7:01 | 28636592 |
| CAGDQAGTALIF | 2 | CAAPQAGTALIF | Melanoma | D008545 | No | BAGE | AARAVFLAL | NA | PBMC | CD8 | NA | TRAV8-2 | TRAJ15 | TRBV12 | NA | TRBJ2-1 | 8921424 |
| CAFVPQGGSEKLVF | 2 | CAVHPQGGSEKLVF | Alzheimer's disease | D000544 | Yes | NA | NA | NA | CSF | CD8 | effector memory CD45RA+ | TRAV13-1 | TRAJ57/58 | TRBV7-9 | NA | TRBJ2-2 | 31915375 |
| CALRTNNNDMRF | 2 | CALLYNNNDMRF | Influenza | D009980 | Yes | Matrix protein (M1) | GILGFVFTL | 20354 | PBMC | CD8 | NA | TRAV19 | NA | TRBV27 | NA | NA | 28636589 |
| CAERMDTGRRALTF | 2 | CAGYMDTGRRALTF | Yellow fever virus | D015005 | Yes | YFV-17D | LLWNGPMAV | 121572 | NA | CD8 | NA | TRAV25 | TRAJ5 | TRBV15 | TRBD2 | TRBJ2-7 | 28103239 |
| CIVRDYKLSF | 2 | CAVRIDYKLSF | Cytomegalovirus (CMV) | D003586 | No | pp65 | NLVPMVATV | 44920 | PBMC | CD8 | NA | TRAV1-4 | TRAJ20 | TRBV6-02 | NA | TRBJ1-1 | 16237109 |
| CIRGIYGGSQGNLIF | 2 | CIRYGGSQGNLIF | Celiac disease | D002446 | Yes | DQ2.5-glia-?2 | NA | NA | PBMC | CD4 | NA | TRAV12-3 | TRAJ22 | NA | NA | NA | 33927715 |

| | | | | | | | | | | | | | | | | | |
|---|---|---|---|---|---|---|---|---|---|---|---|---|---|---|---|---|---|
| CALSDRGGTASKLTF | 2 | CALSENRGGTASKLTF | M. tuberculosis | D009169 | Yes | NA | NA | NA | PBMC | CD8 | NA | NA | TRAJ44 | TRBV20-1 | NA | TRBJ1-305 | 30992377 |
| CAASAFHQTGANNLFF | 2 | CAASAQTGANNLFF | M.Tuberculosis | D009169 | Yes | NA | NA | NA | PBMC | CD4 | NA | TRAV29/DV5 | TRAJ36 | TRBV5-1 | NA | TRBJ2-1 | 32341563 |
| CAVGETGGFKTIF | 2 | CAVNTGGFKTIF | Melanoma | D008545 | No | Melan-A/MART-1 | EAAGIGILTV | 10987 | Tumor Tissue | NA | NA | TRAV2 | TRAJ9 | NA | NA | NA | 7777568 |
| CAFIGGNTPLVF | 2 | CAVRGGNTPLVF | M.Tuberculosis | D009169 | Yes | NA | NA | NA | PBMC | CD4 | NA | TRAV8-1 | TRAJ29 | TRBV3-1 | NA | TRBJ1-1 | 32341563 |
| CAAPNLYSGGGADGLTF | 2 | CAAPYSGGGADGLTF | Influenza | D009980 | Yes | Matrix protein (M1) | GILGFVFTL | 20354 | Bronchoalveolar | CD8 | NA | TRAV12-2:01 | TRAJ4-01 | TRBV2-01 | NA | TRBJ2-5:01 | 28636592 |
| CAAFGGSNYKLTF | 2 | CALGGGSNYKLTF | Influenza | D009980 | Yes | PA | SSLENFRAYV | 61151 | Bronchoalveolar | CD8 | NA | TRAV6D-6:02 | TRAJ5-01 | TRBV2-01 | NA | TRBJ1-4:02 | 28636592 |
| CAERYNQGGKLIF | 2 | CAYRSYNQGGKLIF | Diabetes Type 1 | D003922 | Yes | NA | NA | NA | Pancreatic islets | CD4 | NA | TRAV38-2/DV8 | TRAJ23 | TRBV7-9 | NA | TRBJ2-6 | 27920090 |
| CAVGRPFGNVLHC | 2 | CAVSPFGNVLHC | Influenza | D009980 | Yes | Matrix protein (M1) | GILGFVFTL | 20354 | NA | NA | NA | TRAV14/DV4 | NA | TRBV2 | NA | NA | 30418433 |
| CAYRSENRDDKIIF | 2 | CAYSGNRDDKIIF | M.Tuberculosis | D009169 | Yes | NA | NA | NA | PBMC | CD4 | NA | mTRDV2-2 | TRAJ30 | TRBV19 | NA | TRBJ2-1 | 32341563 |
| CAASGSDGQKLLF | 2 | CAMRGSDGQKLLF | Neoantigen | C050269 | Yes | NSDHL-A9V | KLVALGINAV | NA | NA | NA | NA | TRAV27 | NA | TRBV9 | NA | NA | 30418433 |
| CAENRSGGGNKLTF | 2 | CAENWSGGGGNKLTF | M. tuberculosis | D009169 | Yes | NA | NA | NA | PBMC | CD8 | NA | NA | TRAJ10 | TRBV12-3 | NA | TRBJ2-4 | 30992377 |
| CIVRGTDSWGKLQF | 2 | CIVMTTDSWGKLQF | M.Tuberculosis | D009169 | Yes | NA | NA | NA | PBMC | CD4 | NA | TRAV26-1 | TRAJ24 | TRBV30 | NA | TRBJ1-3 | 32341563 |
| CAVRTNTGNQFYF | 2 | CARNTGNQFYF | Cytomegalovirus (CMV) | D003586 | No | pp65 | NLVPMVATV | 44920 | PBMC | CD8 | NA | TRAV18 | TRAJ49 | TRBV6-03 | NA | TRBJ1-4 | 16237109 |
| CAMNLLQGAQKLVF | 2 | CAASLLQGAQKLVF | Diabetes Type 1 | D003922 | Yes | IGRP | VLFGLGFAI | 103705 | Pancreatic islets | CD8 | NA | TRAV29/DV5:01 | TRAJ54:01 | TRBV3-1:01 | TRBD1:01 | TRBJ2-4:01 | 28300170 |
| CAVPKGDTGRRALTF | 2 | CAVRGDTGRRALTF | M.Tuberculosis | D009169 | Yes | NA | NA | NA | PBMC | CD4 | NA | TRAV8-1 | TRAJ5 | TRBV12-5 | NA | TRBJ1-2 | 32341563 |

| | | | | | | | | | | | | | | | | | |
|---|---|---|---|---|---|---|---|---|---|---|---|---|---|---|---|---|---|
| CAVNNNARLMF | 2 | CAEDNNARLMF | Epstein Barr virus (EBV) | D020031 | No | BMLF-1 | GLCTLVAML | 20788 | Synovial Fluid | CD8 | NA | TRAV15 | TRAJ31 | TRBV20-01 | NA | TRBJ 1-2 | 10925283 |
| CAANDHNNARLMF | 2 | CAASDNNARLMF | M.Tuberculosis | D009169 | Yes | NA | NA | NA | PBMC | CD4 | NA | TRAV29/DV5 | TRAJ31 | TRBV29-1 | NA | TRBJ 1-1 | 32341563 |
| CALTTGANSKLTF | 2 | CALIAGANSKLTF | Influenza | D009980 | Yes | Matrix protein (M1) | GILGFVFTL | 20354 | PBMC | CD8 | NA | TRAV9-2:01 | TRAJ5-01 | TRBV19:02 | NA | TRBJ 2-3:01 | 28636592 |
| CAVEYGGSQGNLIF | 2 | CIVYGGSQGNLIF | Human immunodeficiency virus (HIV) | D006678 | No | RF10 Protein Nef | RYPLTFGWCF | 56620 | PBMC | CD8 | NA | TRAV26-1 | TRAJ42 | TRBV23-01 | TRBD1 | TRBJ 1-5 | 24899498 |
| CAVYTGGGNKLTF | 2 | CAEVSTGGGNKLTF | Human immunodeficiency virus (HIV) | D006678 | No | RF10 Protein Nef | RYPLTFGWCF | 56620 | PBMC | CD8 | NA | TRAV13-2 | TRAJ10 | TRBV4-01 | TRBD1 | TRBJ 1-1 | 24899498 |
| CAVGTNAGKSTF | 2 | CAVNAGKSTF | Yellow fever virus | D015005 | Yes | YFV-17D | LLWNGPMAV | 121572 | NA | CD8 | NA | TRAV12-2 | TRAJ27 | TRBV6-06 | TRBD1, TRBD2 | TRBJ 2-7 | 28103239 |
| CAVRAARLMF | 2 | CAVTSARLMF | Yellow fever virus | D015005 | Yes | YFV-17D | LLWNGPMAV | 121572 | NA | CD8 | NA | TRAV12-2 | TRAJ31 | TRBV4-03 | TRBD1 | TRBJ 2-7 | 28103239 |
| CAYGVNRDDKIIF | 2 | CAGVGRDDKIIF | Breast Cancer | D001943 | No | NA | NA | NA | Breast | CD8 | TIL | TRAV25 | TRAJ30 | TRBV7-02 | NA | TRBJ 1-5 | 27307436 |
| **TRAV1-2 negative CDR3beta sequence similarity** | | | | | | | | | | | | | | | | | |
| Query | Dist | CDR3.beta.aa | Pathology | Pathology.Mesh.ID | Single.cell | Antigen.protein | Epitope.peptide | Epitope.ID | Tissue | T.Cell.Type | T.cell.characteristics | TRAV | TRAJ | TRBV | TRBD | TRBJ | PubMed.ID |
| CATSRDLPGNSPLHF | 2 | CATSRDRGGNSPLHF | M.Tuberculosis | D009169 | Yes | NA | NA | NA | PBMC | CD4 | NA | TRAV9-1 | TRAJ7 | TRBV15 | NA | TRBJ 1-6 | 32341563 |
| CSARDGTSGDTGELFF | 1 | CSARDGTSDTGELFF | M. tuberculosis | D009169 | Yes | NA | NA | NA | PBMC | CD8 | NA | NA | TRAJ33 | TRBV20-1 | NA | TRBJ 2-2 | 30992377 |
| CASSLRPSNQPQHF | 2 | CASSLGPGNQPQHF | M.Tuberculosis | D009169 | Yes | NA | NA | NA | PBMC | CD4 | NA | mTRDV2-2 | TRAJ40 | TRBV11-1 | NA | TRBJ 1-5 | 32341563 |
| CASSLTGGLGTEAFF | 2 | CASSLDGGGGTEAFF | M.Tuberculosis | D009169 | Yes | NA | NA | NA | PBMC | CD4 | NA | TRAV21 | TRAJ15 | TRBV5-1 | NA | TRBJ 1-1 | 32341563 |
| CSARQGATEAFF | 2 | CSARPGDTEAFF | M. tuberculosis | D009169 | Yes | NA | NA | NA | PBMC | CD8 | MAIT | NA | TRAJ33 | TRBV20-1 | NA | TRBJ 1-1 | 30992377 |

| | | | | | | | | | | | | | | | | | |
|---|---|---|---|---|---|---|---|---|---|---|---|---|---|---|---|---|---|
| CASSQDGSTDTQYF | 2 | CASSAPGSTDTQYF | M. tuberculosis | D009169 | Yes | NA | NA | NA | PBMC | CD8 | iNKT | TRAV3 | TRAJ18 | TRBV25-1 | NA | TRBJ 2-3 | 30992377 |
| CASSYSNSGSGANVLTF | 2 | CASSYSGGSGANVLTF | M. tuberculosis | D009169 | Yes | NA | NA | NA | PBMC | CD8 | NA | NA | TRAJ33 | TRBV6-2 | NA | TRBJ 2-6 | 30992377 |
| CASSELAGGQETQYF | 2 | CASLLAGGQETQYF | M.Tuberculosis | D009169 | Yes | NA | NA | NA | PBMC | CD4 | NA | TRAV26-1 | TRAJ39 | TRBV2 | NA | TRBJ 2-5 | 32341563 |
| CASSPRPLYEQYF | 2 | CASSIRPSYEQYF | Influenza | D009980 | Yes | Matrix protein (M1) | GILGFVFTL | 20354 | PBMC | CD8 | NA | TRAV8-2:01 | TRAJ41:01 | TRBV19:01 | TRBD2:01 | TRBJ 2-7:01 | 28300170 |
| CASSPGLADIDTQYF | 2 | CASSLGLAGIDTQYF | M.Tuberculosis | D009169 | Yes | NA | NA | NA | PBMC | CD4 | NA | TRAV19 | TRAJ53 | TRBV5-1 | NA | TRBJ 2-3 | 32341563 |
| CASSEQGNTGELFF | 2 | CASSAGNTGELFF | M. tuberculosis | D009169 | Yes | NA | NA | NA | PBMC | CD8 | NA | NA | TRAJ20 | TRBV6-4 | NA | TRBJ 2-2 | 30992377 |
| CASSRSQGNTEAFF | 2 | CASSPSQGVTEAFF | M.Tuberculosis | D009169 | Yes | NA | NA | NA | PBMC | CD4 | NA | TRAV12-3 | TRAJ40 | TRBV18 | NA | TRBJ 1-1 | 32341563 |
| CASSVTGYEQFF | 2 | CASSHSGYEQFF | Yellow fever virus | D015005 | Yes | YFV-17D | LLWNGPMAV | 121572 | NA | CD8 | NA | TRAV12-1 | TRAJ50 | TRBV4-01 | TRBD1, TRBD2 | TRBJ 2-1 | 28103239 |
| CASSLTGEYNEQFF | 2 | CARSTGEYNEQFF | M. tuberculosis | D009169 | Yes | NA | NA | NA | PBMC | CD8 | NA | NA | TRAJ33 | TRBV30 | NA | TRBJ 2-1 | 30992377 |
| CASSLRHLNTEAFF | 2 | CASSLHMNTEAFF | M.Tuberculosis | D009169 | Yes | NA | NA | NA | PBMC | CD4 | NA | TRAV3 | TRAJ15 | TRBV7-2 | NA | TRBJ 1-1 | 32341563 |
| CASSLDGTSGVTDTQYF | 2 | CASSDGTSGGTDTQYF | M. tuberculosis | D009169 | Yes | NA | NA | NA | PBMC | CD8 | NA | NA | TRAJ20 | TRBV6-4 | TRBD2 | TRBJ 2-3 | 30992377 |
| CASRNGGYEQYF | 2 | CASGDGGYEQYF | Influenza | D009980 | Yes | NP | ASNENMETM | 4602 | Bronchoalveolar | CD8 | NA | TRAV14D-3/DV8:08 | TRAJ2-01 | TRBV1-01 | NA | TRBJ 2-7:01 | 28636592 |
| CASSPLGTGNNEQYF | 2 | CASSFLGTGLNEQYF | Hepatitis C virus (HCV) | D006526 | Yes | HCV-KLV(PE) | KLVALGINAV | 32208 | NA | NA | NA | TRAV19 | NA | TRBV28 | NA | NA | 30418433 |
| CASSEASGGADTQYF | 2 | CASSDSGGADTQYF | M. tuberculosis | D009169 | Yes | NA | NA | NA | PBMC | CD8 | NA | NA | TRAJ9 | TRBV6-4 | NA | TRBJ 2-3 | 30992377 |
| CASSESGGGETQYF | 2 | CASSDASGGGETQYF | M. tuberculosis | D009169 | Yes | NA | NA | NA | PBMC | CD8 | NA | NA | TRAJ20 | TRBV6-1 | NA | TRBJ 2-5 | 30992377 |
| CASSLGSRNEQFF | 2 | CASSILGSYNEQFF | Tumor associated antigen (TAA) | D018290 | Yes | EphA2 | IMNDMPIYM | NA | PBMC | CD8 | NA | TRAV19 | TRAJ23 | TRBV19 | NA | TRBJ 2-1 | 33562731 |

| | | | | | | | | | | | | | | | | | |
|---|---|---|---|---|---|---|---|---|---|---|---|---|---|---|---|---|---|
| CASSLGGLDGYTF | 2 | CASSDGGKDGYTF | Hepatitis C virus (HCV) | D006526 | Yes | HCV-KLV | KLVALGINAV | 32208 | NA | NA | NA | NA | NA | TRBV25-1 | NA | NA | 30418433 |
| CASSQVSGNTEAFF | 2 | CASSLVSENTEAFF | M.Tuberculosis | D009169 | Yes | NA | NA | NA | PBMC | CD4 | NA | TRAV8-6 | TRAJ36 | TRBV5-1 | NA | TRBJ1-1 | 32341563 |
| CASSVRLSTDTQYF | 2 | CASSARSTDTQYF | Influenza | D009980 | Yes | Matrix protein (M1) | GILGFVFTL | 20354 | PBMC | CD8 | NA | TRAV6-01 | TRAJ4-01 | TRBV19:02 | NA | TRBJ2-3:01 | 28636592 |
| CASSDGQGREKLFF | 2 | CASSAGQGGEKLFF | M.Tuberculosis | D009169 | Yes | NA | NA | NA | PBMC | CD4 | NA | TRAV10 | TRAJ18 | TRBV25-1 | NA | TRBJ1-4 | 32341563 |
| CASSSGTGDTGELFF | 2 | CASSAGTGHTGELFF | M. tuberculosis | D009169 | Yes | NA | NA | NA | PBMC | CD8 | NA | NA | TRAJ12 | TRBV6-1 | NA | TRBJ2-2 | 30992377 |
| CASKLGQGGYEQYF | 2 | CASSFGQGGYEQYF | Influenza | D009980 | Yes | Matrix protein (M1) | GILGFVFTL | 20354 | PBMC | CD8 | NA | TRAV8-2 | NA | TRBV13 | NA | NA | 28636589 |
| CASSFGDRGREQYF | 2 | CASSQDRGREQYF | M.Tuberculosis | D009169 | Yes | NA | NA | NA | PBMC | CD4 | NA | TRAV17 | TRAJ8 | TRBV4-3 | NA | TRBJ2-7 | 32341563 |
| CASSQAAGGNTDTQYF | 2 | CASSEAAGTGNTDTQYF | M. tuberculosis | D009169 | Yes | NA | NA | NA | PBMC | CD8 | NA | NA | TRAJ33 | TRBV6-1 | NA | TRBJ2-3 | 30992377 |
| CASSSRQGHTGELFF | 2 | CASSDRGHTGELFF | M. tuberculosis | D009169 | Yes | NA | NA | NA | PBMC | CD8 | NA | NA | TRAJ49 | TRBV6-4 | NA | TRBJ2-2 | 30992377 |
| CASSEMGTGELFF | 2 | CASSAQGTGELFF | M.Tuberculosis | D009169 | Yes | NA | NA | NA | PBMC | CD4 | NA | TRAV21 | TRAJ56 | TRBV3-1 | NA | TRBJ2-2 | 32341563 |
| CASSDGSGGYGYTF | 2 | CASSDGTGPYGYTF | Diabetes Type 1 | D003922 | Yes | NA | NA | NA | Pancreatic islets | CD4 | NA | TRAV22 | TRAJ28 | TRBV6-1 | NA | TRBJ1-2 | 27920090 |
| CASSLGLAGTTDTQYF | 2 | CASSGLAGGTDTQYF | M. tuberculosis | D009169 | Yes | NA | NA | NA | PBMC | CD8 | MAIT | NA | TRAJ33 | TRBV6-4 | TRBD2 | TRBJ2-3 | 30992377 |
| CASSQHDGYTQYF | 2 | CASSQVDGYEQYF | M.Tuberculosis | D009169 | Yes | NA | NA | NA | PBMC | CD4 | NA | TRAV9-1 | TRAJ44 | TRBV3-1 | NA | TRBJ2-7 | 32341563 |
| CASSTGQGNTIYF | 2 | CASSAGGGNTIYF | M.Tuberculosis | D009169 | Yes | NA | NA | NA | PBMC | CD4 | NA | TRAV13-1 | TRAJ45 | TRBV5-1 | NA | TRBJ1-3 | 32341563 |
| CASSDSSGGANEQFF | 2 | CASSDASGGAYNEQFF | M. tuberculosis | D009169 | Yes | NA | NA | NA | PBMC | CD8 | NA | NA | TRAJ33 | TRBV6-4 | NA | TRBJ2-1 | 30992377 |
| CASSDPERTEAFF | 2 | CASSDAENTEAFF | Cytomegalovirus (CMV) | D003586 | Yes | pp65 | NLVPMVATV | 44920 | PBMC | CD8 | NA | TRAV1-2:01 | TRAJ3-01 | TRBV6-01 | NA | TRBJ1-1:01 | 28636592 |

| | | | | | | | | | | | | | | | | | |
|---|---|---|---|---|---|---|---|---|---|---|---|---|---|---|---|---|---|
| CASSPGTGLSYEQYF | 2 | CASSEGTGGSYEQYF | M. tuberculosis | D009169 | Yes | NA | NA | NA | PBMC | CD8 | MAIT | NA | TRAJ33 | TRBV6-1 | TRBD1 | TRBJ 2-7 | 30992377 |
| CASSAPLNNEQFF | 2 | CASSHPLGNEQFF | M.Tuberculosis | D009169 | Yes | NA | NA | NA | PBMC | CD4 | NA | TRAV8-1 | TRAJ31 | TRBV4-1 | NA | TRBJ 2-1 | 32341563 |
| CASSLQGQGADEQFF | 2 | CASSLEGQGASEQFF | Celiac disease | D002503 | Yes | DQ2-a-I | PFPQPELPY | 47539 | Small intestine | CD4 | NA | TRAV9-1 | TRAJ33-1 | TRBV5-5*01 | TRBD2*01 | TRBJ 2-1 | 24777060 |
| CASSLNGGYNEQFF | 2 | CASSALAGGYNEQFF | Neoantigen | C050269 | Yes | KIF20B-YTS-S6L | YTSEILSPI | NA | NA | NA | NA | NA | NA | TRBV2 | NA | NA | 30418433 |
| CASSSMLSETQYF | 2 | CASSQLSETQYF | M.Tuberculosis | D009169 | Yes | NA | NA | NA | PBMC | CD4 | NA | TRAV12-1 | TRAJ45 | TRBV3-1 | NA | TRBJ 2-5 | 32341563 |
| CSARGLGNTEAFF | 2 | CSARDLGNLEAFF | M.Tuberculosis | D009169 | Yes | NA | NA | NA | PBMC | CD4 | NA | TRAV9-1 | TRAJ23 | TRBV20-1 | NA | TRBJ 1-1 | 32341563 |
| CASSHRNTGELFF | 2 | CASSAGNTGELFF | M. tuberculosis | D009169 | Yes | NA | NA | NA | PBMC | CD8 | NA | NA | TRAJ20 | TRBV6-4 | NA | TRBJ 2-2 | 30992377 |
| CASSLELLVEQFF | 2 | CASSLTLLEQFF | M.Tuberculosis | D009169 | Yes | NA | NA | NA | PBMC | CD4 | NA | TRAV26-1 | TRAJ54 | TRBV13 | NA | TRBJ 2-1 | 32341563 |
| CSASLSDYGYTF | 2 | CSASRTDYGYTF | M.Tuberculosis | D009169 | Yes | NA | NA | NA | PBMC | CD4 | NA | mTRAV14D-1 | TRAJ8 | TRBV20-1 | NA | TRBJ 1-2 | 32341563 |
| CATSRDKGVNQPQHF | 2 | CATSRDPGPNQPQHF | M.Tuberculosis | D009169 | Yes | NA | NA | NA | PBMC | CD4 | NA | TRAV26-1 | TRAJ24 | TRBV15 | NA | TRBJ 1-5 | 32341563 |
| CASSLAMGLYEQYF | 2 | CASKSLAGGLYEQYF | M.Tuberculosis | D009169 | Yes | NA | NA | NA | PBMC | CD4 | NA | TRAV17 | TRAJ57 | TRBV2 | NA | TRBJ 2-7 | 32341563 |
| CASSLDPSDSGANVLTF | 2 | CASSLDGDSGANVLTF | M. tuberculosis | D009169 | Yes | NA | NA | NA | PBMC | CD8 | NA | NA | TRAJ33 | TRBV6-2 | NA | TRBJ 2-6 | 30992377 |
| CASSEDPTDTQYF | 2 | CASSEGITDTQYF | M.Tuberculosis | D009169 | Yes | NA | NA | NA | PBMC | CD4 | NA | TRAV13-1 | TRAJ33 | TRBV7-2 | NA | TRBJ 2-3 | 32341563 |
| CASSPQHPQETQYF | 2 | CASSPWQHQETQYF | M.Tuberculosis | D009169 | Yes | NA | NA | NA | PBMC | CD4 | NA | TRAV21 | TRAJ13 | TRBV18 | NA | TRBJ 2-5 | 32341563 |
| CASSFGSDSNQPQHF | 2 | CASSFADSNQPQHF | M.Tuberculosis | D009169 | Yes | NA | NA | NA | PBMC | CD4 | NA | TRAV25 | TRAJ22 | TRBV11-2 | NA | TRBJ 1-5 | 32341563 |
| CASSQTGNQPQHF | 2 | CASQGNQPQHF | M.Tuberculosis | D009169 | Yes | NA | NA | NA | PBMC | CD4 | NA | TRAV26-1 | TRAJ5 | TRBV28 | NA | TRBJ 1-5 | 32341563 |
| CASSRGGGSEKLFF | 2 | CASSGRGGGEKLFF | M. tuberculosis | D009169 | Yes | NA | NA | NA | PBMC | CD8 | iNKT | NA | TRAJ18 | TRBV25-1 | NA | TRBJ 1-4 | 30992377 |

| | | | | | | | | | | | | | | | | | |
|---|---|---|---|---|---|---|---|---|---|---|---|---|---|---|---|---|---|
| CASSFGGGRSSYNEQFF | 2 | CASSDSGGRSSYNEQFF | M.Tuberculosis | D009169 | Yes | NA | NA | NA | PBMC | CD4 | NA | TRAV10 | TRAJ18 | TRBV25-1 | NA | TRBJ2-1 | 32341563 |
| CASSGTGRVDTQYF | 2 | CASSGTGQDTQYF | mCMV | D018146 | Yes | m139 | TVYGFCLL | 67227 | Spleen | CD8 | NA | TRAV7-2:02 | TRAJ5-01 | TRBV4-01 | NA | TRBJ2-5:01 | 28636592 |
| CASSLGATGRASGNTIYF | 2 | CASSLGHTGASGNTIYF | M.Tuberculosis | D009169 | Yes | NA | NA | NA | PBMC | CD4 | NA | TRAV5 | TRAJ34 | TRBV5-1 | NA | TRBJ1-3 | 32341563 |
| CASSESSTTNEKLFF | 2 | CASSESETGNEKLFF | M.Tuberculosis | D009169 | Yes | NA | NA | NA | PBMC | CD4 | NA | TRAV10 | TRAJ18 | TRBV25-1 | NA | TRBJ1-4 | 32341563 |
| CASSLVASGGANEQFF | 2 | CASSGASGGANEQFF | M. tuberculosis | D009169 | Yes | NA | NA | NA | PBMC | CD8 | NA | NA | TRAJ33 | TRBV6-1 | NA | TRBJ2-1 | 30992377 |
| CASSYDRPNTEAFF | 2 | CASSDRGNTEAFF | M.Tuberculosis | D009169 | Yes | NA | NA | NA | PBMC | CD4 | NA | TRAV8-4 | TRAJ40 | TRBV6-1 | NA | TRBJ1-1 | 32341563 |
| CASSLLGTETQYF | 2 | CASSLAGTDTQYF | M.Tuberculosis | D009169 | Yes | NA | NA | NA | PBMC | CD4 | NA | TRAV12-2 | TRAJ54 | TRBV5-6 | NA | TRBJ2-3 | 32341563 |
| CASSFGGGLGNTEAFF | 2 | CASSSGGGLNTEAFF | M.Tuberculosis | D009169 | Yes | NA | NA | NA | PBMC | CD4 | NA | TRAV19 | TRAJ39 | TRBV7-2 | NA | TRBJ1-1 | 32341563 |
| CASSQDGTSGYNEQFF | 2 | CASSDGTSGNEQFF | M. tuberculosis | D009169 | Yes | NA | NA | NA | PBMC | CD8 | MAIT | NA | TRAJ33 | TRBV6-4 | NA | TRBJ2-1 | 30992377 |
| CASSLGQLNQPQHF | 2 | CASSEGLNQPQHF | M.Tuberculosis | D009169 | Yes | NA | NA | NA | PBMC | CD4 | NA | mTRAV14D-1 | TRAJ15 | TRBV6-1 | NA | TRBJ1-5 | 32341563 |
